# Reduced Dietary Protein Induces Changes in the Dental Proteome

**DOI:** 10.1101/2025.01.13.632248

**Authors:** Robert W. Burroughs, Christopher J. Percival, Natasha S. Vitek

## Abstract

Experimental studies have demonstrated that nutritional changes during development can result in phenotypic changes to mammalian cheek teeth. This developmental plasticity of tooth morphology is an example of phenotypic plasticity. Because tooth development occurs through complex interactions between manifold processes, there are many potential mechanisms which can contribute to a tooth’s norm of reaction. Determining the identity of those mechanisms and the relative importance of each of them is one of the main challenges to understanding phenotypic plasticity. Quantitative proteomics combined with experimental studies allow for the identification of potential molecular contributors to a plastic response through quantification of expressed gene products. Here, we present the results of a quantitative proteomics analysis of mature upper first molars in *Mus musculus* from a controlled feeding experiment. Pregnant and nursing mothers were fed either a low-dietary protein (10%) treatment diet or control (20%) diet. Low-dietary protein was not associated with reduced molar size or skull length. However, expression of tooth-related proteins, immune system proteins, and actin-based myosin proteins were significantly altered in our low-dietary protein proteomics sample. The differential expression of immune proteins along with systematic reduction in actin-based myosin protein expression are novel discoveries for tooth proteomics studies. We propose that studies that aim to elucidate specific mechanisms of molar phenotypic plasticity should prioritize investigations into the relationships between IGF regulation and tooth development and actin-based myosin expression and tooth development.

**Research Highlights:** A low-protein diet during development results in significantly altered protein expression for odontogenetic and osteogenic proteins, immune system proteins, and actin-based myosin proteins within *Mus musculus*, but does not alter skull length or molar size.

Graphical Abstract

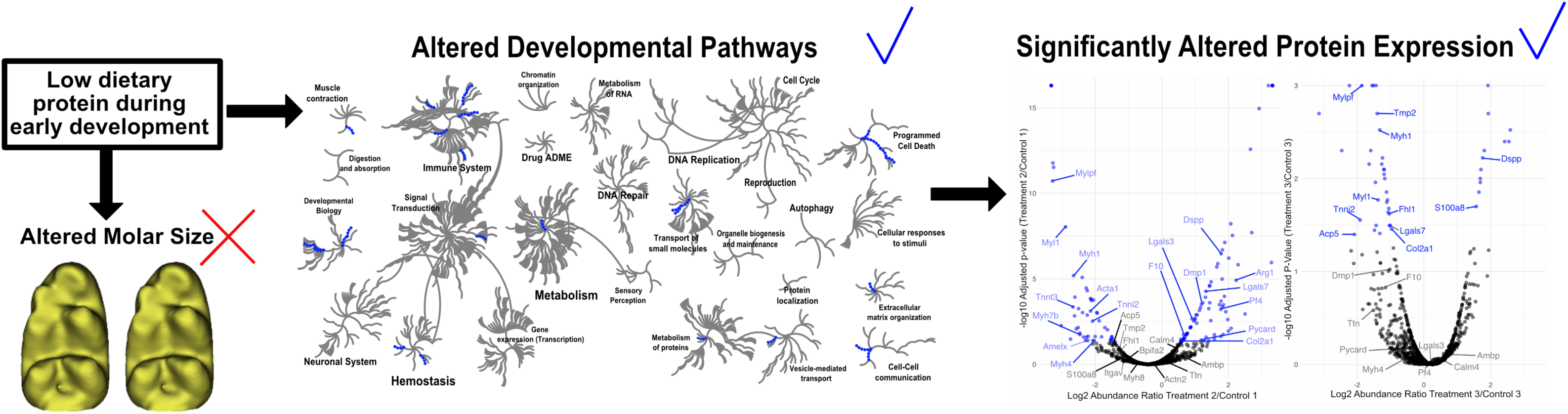

## Introduction

Phenotypic plasticity, the differential expression of a phenotype, is often invoked as a way that organisms respond to changing environments. Within lab and common garden experiments, nutritional changes such as changes in the quantity of dietary protein induce plastic changes to mammalian cheek tooth phenotypes (Patton & Brylski, 1987; Paynter & Grainer, 1956; Shaw & Griffiths, 1963). Typically, these are changes in tooth size, size and shape of tooth cusps, and timing of eruption (Holloway et al., 1961; Patton & Brylski, 1987; Paynter & Grainer, 1956; Searle, 1954; Shaw & Griffiths, 1963). This demonstrates that there is plasticity in dental development that can provide a short-term, non-evolutionary response to changing environments (Levis & Pfennig, 2021).

However, measures of plastic phenotypic response to environmental change often do not identify the molecular and genetic pathways underlying those changes. The use of quantitative proteomics to quantify differences in protein expression could allow for the identification and study of pathways that are altered by an environmental change (e.g., are plastic). Here, we present a study characterizing protein expression variation in a sample of upper first molars (M1s) and phenotypic variation in lower first molars (lower M1s) from lab mice fed a low-protein diet as part of a controlled feeding experiment. This experiment was designed to induce developmental changes to dental traits and to identify proteins and associated pathways that may underlie phenotypic plasticity. This allows us to characterize those pathways most impacted by exposure to low dietary protein during embryonic and early postnatal tooth development.

### Proteomics

Proteomics is broadly the field focused on identifying, annotating, and quantifying variation of proteins. Quantification of variation includes both protein sequence variation and protein expression variation. Informatics approaches are applied to protein spectral data collected via tandem liquid-chromatography mass-spectrometry (LC-MS/MS) (Heck & Neely, 2020). Protein expression profiles are typically tissue– and developmental stage-dependent and must be interpreted within the specific spatiotemporal contexts in which samples were collected (Rebeaud et al., 2021).

An assemblage of proteins that is expressed within a specific tissue or structure is often referred to as a ‘proteome’ (Sharma et al., 2020). For example, within the dental proteome enamel and dentin forming proteins are found in high abundance (Sharma et al., 2020). However, outside of dental tissues, these proteins are detected in low abundance and only at certain developmental stages (Bansal et al., 2012; Ritchie, 2018).

If basic processes of tooth mineralization are impacted by dietary deficiency, we expect a change in associated protein expression and a change in phenotype such as occlusal pattern, size, and/or shape (Harjunmaa et al., 2014). It is possible that some aspects of tooth mineralization are more easily perturbed by environmental change than others and their associated pathways may more frequently underlie plastic changes. Measuring the tooth proteome may allow us to identify potential candidates underlying plastic changes in tooth phenotype.

Beyond stereotypical tooth-associated proteins, previous characterizations of the dental proteome have suggested that teeth serve as reservoirs of more general patterns of organismal protein expression during amelogenesis (tooth mineralization) (Froment et al., 2021; Giovani et al., 2021; Green et al.; Sharma et al., 2020). For example, Green et al. (2019) recovered immune system related proteins from enamel that was micro sampled from near the enamel-dentin junction of mineralizing pig molars. These immune proteins currently have no known function in amelogenesis. These proteins are definitively present within enamel tissue, but they may originate in nearby salivary glands that are in contact with enamel cell proliferation zones and regions of mineralization at the enamel-dentin junction (Green et al., 2019; Jagr et al., 2019). If mineralized teeth serve as an archive of broader organismal protein expression at the time of mineralization, it is possible that tooth proteomic data could provide evidence of more generalized responses to environmental perturbations.

### This Study

To determine what pathways are altered by an impoverished diet, we conducted a controlled feeding experiment and investigated protein expression in the cheek teeth. The study presented here is part of a larger research program that will also investigate the impacts of low dietary nutrition on a variety of tooth phenotypes. Pregnant and nursing mouse dams were fed a low protein or control protein diet and effects were measured in their offspring. Previous feeding studies on dietary protein quantity in rats and mice suggested that a threshold of 10-12% (by weight) dietary protein reduction from a control of 20-24% should induce phenotypically plastic changes to molar size, skull length, and disrupt timing of molar development (Barbeito-Andrés et al., 2016 2016; Holloway et al., 1961; Miller & German, 1999; Paynter & Grainer, 1956; Paynter & Grainer, 1961; Pucciarelli, 1980; Shaw & Griffiths, 1963).

With this context we anticipated that reduction in dietary protein would disrupt normal tooth development and alter protein expression patterns during embryonic and early postnatal development. We predicted a significant reduction in M1 size, consistent with previous reports (e.g., (Holloway et al., 1961; Paynter & Grainer, 1956; Paynter & Grainer, 1961; Shaw & Griffiths, 1963) and significant reduction in skull length based on previous studies in rat and mouse (e.g., (Barbeito-Andrés et al., 2016; Lobe et al., 2006; Miller & German, 1999; Pucciarelli, 1980). Previous studies suggested that halving dietary protein would likely increase risk of infection and increase metabolic stress in low protein mice (Giovani et al., 2021; Steward et al., 2023). Thus, we anticipated that there might be additional proteomic signals related to stress or immune system function if those protein signatures are preserved within dentition. Recovery of significant differential expression of proteins because of a dietary change allowed us to identify and rank the molecular pathways most likely to be perturbed by environmental influences like dietary protein reduction.

## Methods

### Data Availability

All datasets, code, workflows, and raw phenotypic data reported are available as Supplemental Materials on Dryad: Dryad DOI. Raw proteomics data are available on MassIVE database by querying the following MassIVE identifier: MSV000098600.

### Feeding Experiment

A breeding colony of inbred strain C57BL/6J (RRID: IMSR JAX:000664) mice was established at the Division of Laboratory Animal Research at Stony Brook University, in accordance with authorized IACUC protocol (SBU IACUC 2023-0014). Male and female mice were acquired at 8 weeks of age and housed in residence to acclimate until breeding began at 12 weeks of age. Males were placed with females for up to 72 hours (3 day-night cycles) and females were checked daily for presence of a copulatory plug. Once a plug was present or 72 hours had passed, the male and female were separated. Females, regardless of plug presence, were then randomly assigned to control or treatment diets to ensure that any developing embryos were on as consistent a diet as possible. Mice assigned to the control group were fed a 20% raw protein diet (PicoLab Rodent Diet 20, 5053). Mice assigned to treatment were fed a 10% raw protein diet (Mod LabDiet 5053 with ∼10% Protein Red, 5BQM). Prior to this assignment, all specimens consumed the control diet.

If females were not pregnant, as evidenced by swollen abdomen after ∼7 days post-mating attempt, low-protein females were cycled back to the control protein (20%) diet. They were re-acclimated to that diet for 14 days before additional mating was attempted. To ensure that additional stress was not induced via single housing, pairs of females were housed together during acclimation and pregnancy. Mating timings were staggered to ensure that two females in the same cage would not give birth at the same time. This allowed for the separation of females and their litters once they had given birth.

Pregnant and nursing females in the treatment group were fed only the low-protein diet for the remainder of their lifespan. Offspring were weaned at approximately ∼21 days postnatal (P21). Siblings were housed with their respective sex and fed their respective diets until P28 to ensure full eruption of the third molars. Dams and offspring were euthanized when the offspring were P28. Dams were not reused to avoid introducing bias related to improvement of maternal care from first to second litter (Weber & Olsson, 2008). A total of 12 litters (7 treatment, 5 control) constituting 74 offspring (treatment n=34, control n=30) from this feeding experiment were collected. Aside from the 74 collected offspring, there was postnatal attrition of one full litter of treatment pups (n=8), where the mother declined to nurse the pups. The smallest pup from each of five litters (treatment n=3; control n=2) also did not survive to weaning.

### Sampling

For proteomics data we opportunistically sampled specimens from four litters of our feeding experiment. In our sample, males were the more common sex, and thus we selected by random, four male sibling pairs for destructive sampling. Selecting only males also removed variation in protein expression that would be associated with specimen sex within our small sample. The upper first molar (upper M1) was selected for protein extraction because of its ease of extraction and because it and its lower jaw counterpart are the first molars to develop and erupt (Figure 1A). Eight upper M1s (treatment n=4; control n=4) were extracted from sibling pairs from four litters immediately after euthanasia by dissecting away the maxillary gingiva and exposed maxillary bone to reveal tooth roots. A blunt probe was used to lever the tooth out. Excess tissue, including any root bundle, was removed with forceps. Extracted teeth were then washed with 70% ETOH, wiped dry with a clean Kimwipe, and stored in new cryotubes. Teeth were placed into a –80°C freezer and maintained at –80°C until preparation for protein quantification.

**Figure 1:**
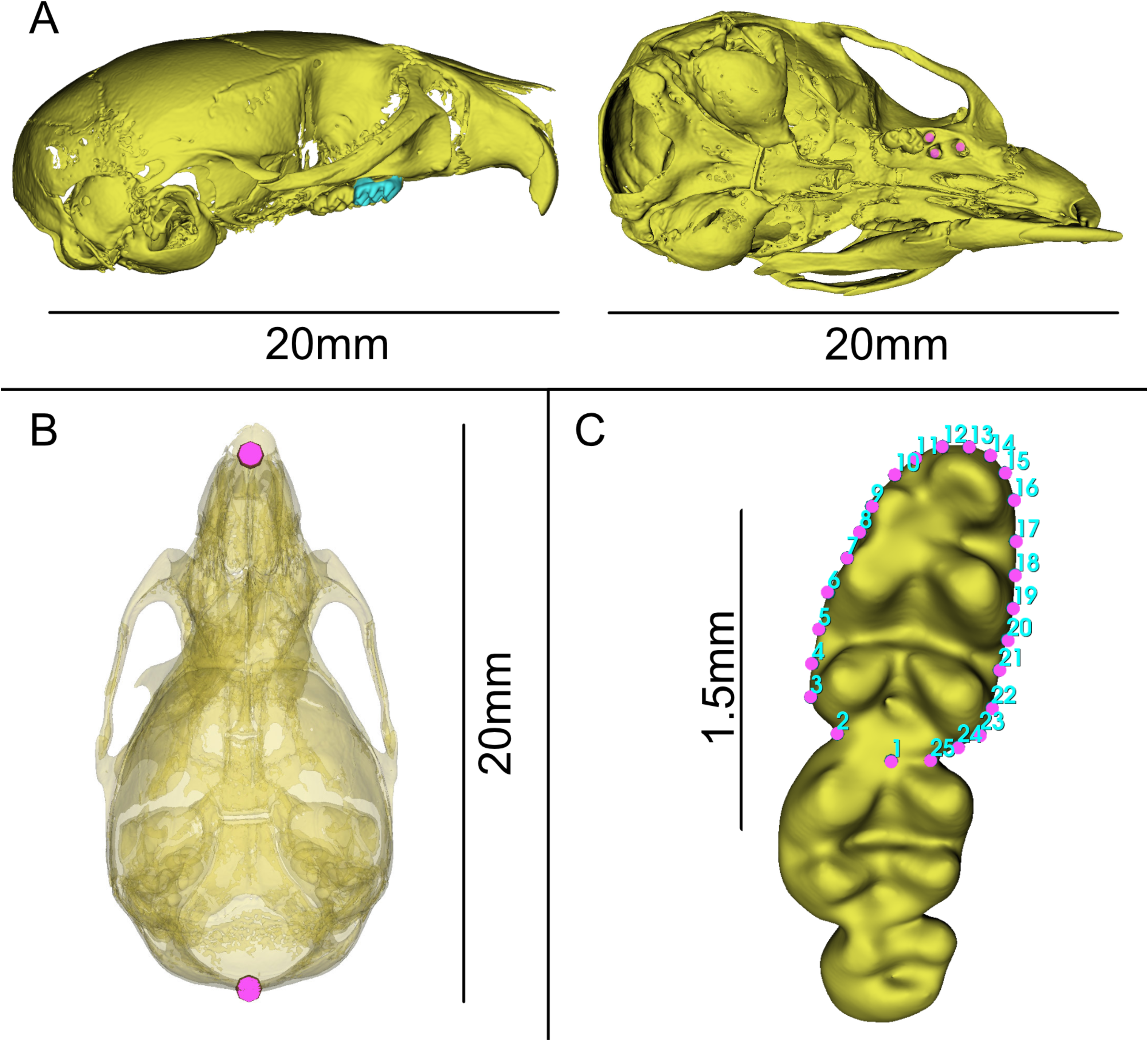
Models of skulls of treatment specimens used for proteomic sampling and processed from µCT scans. A: Lateral and Ventral view of skull, showing where right upper M1 was removed, M2 and M3 are still visible. Light blue shows M1 still in place, Fuschia dots indicate where roots for M1 once inserted. B: Dorsal view of skull, opacity adjusted to 50% to show location of landmarks for straight-line skull estimate. C: Scheme of 25 landmarks placed on left lower m1 used to assess crown area and shape between treatment and control specimens.

For phenotypic data, we sampled the left lower first molar (lower M1) of specimens from four treatment and four control litters. These litters include the four litters sampled for proteomics. We measured the phenotype of 50 (treatment n=24; control n=26), where 24 were males and 26 were females. Three of the specimens sampled for proteomics (2 controls, 1 treatment) were excluded from the phenotypic data because they were held out for future synchrotron scanning. Specimen data are included in Supplemental Materials.

### Phenotyping

We utilized µCT imaging to generate high-fidelity 3D surfaces of mouse specimens. This approach allows us to utilize semi-automated phenotyping approaches. We batch scanned specimens at the New York Institute of Technology Visualization Center on a Bruker Skyscan 1173 at 75kV, 106µA, at a resolution of 19.938µm. Specimens were decapitated post-euthanasia and ethafoam was inserted to separate the upper and lower dentition. Heads were then wrapped in Kimwipes and stacked vertically in 50ml Falcon tubes, with males and females grouped separately within the tube and one to two litters of the same type (e.g., all control or all treatment) per tube. Litters were separated by stacked ethafoam discs that form a visual barrier in reconstructed scan data.

Post-processing of µCT data was done at Stony Brook University, by author RWB, utilizing 3D Slicer (Fedorov et al., 2012). Reconstructed batch scans were loaded into Slicer and individual specimens were cropped from the batch scan volume. Minimum thresholding was used to segment all bony elements within the scan. The Islands segmentation tool was used to separate out the largest element within the segment, the cranium. Once segmented, crania were converted to 3D models. It was necessary to segment the entire tooth row, as opposed to isolate the lower M1, due to contact between teeth. Minimum thresholding was again applied, but with increased minima compared to bony elements to isolate the denser enamel. The Scissors tool was used to circle the lower left tooth row and “Erase Outside” was selected to remove all remaining elements within the segment. Finally, the Islands tool was again used to keep only the largest element within the segment (the tooth row). Once segmented, toothrows were converted to 3D models.

We utilized landmarks to measure phenotypic data on 3D models. To place landmarks, we utilized Automated Landmarking through Point cloud Alignment and Correspondence Analysis (ALPACA) implemented within the 3D Slicer SlicerMorph extension (Porto et al., 2021; Rolfe et al., 2021). The strength of ALPACA lies in rapid implementation and reduced user bias in landmark placement for landmark projection. ALPACA accomplishes this with a point cloud-based deformation of a reference landmark template and mesh to match non-reference specimens. Once the deformation has occurred, landmarks from the reference specimen are projected onto the non-reference specimen. Because this method utilizes a deformation-based approach it is necessary that specimens be similar in shape (as they are in this intraspecific study of a single tooth) and that placed landmarks be quality checked after running ALPACA. We generated two reference templates: 1) a control cranium with two landmarks for calculating straight-line skull length, 2) a control lower M1 with 25 landmarks representing tooth outline/crown footprint (Figures 1B and 1C). After ALPACA was run, a single author (RWB) checked all of ALPACAs landmark position predictions and moved any landmarks that were not properly projected.

Analysis of landmark data was conducted in **R** with core R tools (RCoreTeam, 2023), the **geomorph** (Adams et al., 2013) package, with visualization via the core **plot** functions and **ggplot** package (Wickham, 2016). The landmark files and R script are in Supplemental Materials. To test hypotheses of changes in skull-length we measured the line between landmarks at the apex of the foramen magnum and the junction between the maxillae (Figure 1B). A Welch’s T-Test was calculated using the **t.test** function in R to determine if there was a significant difference between treatment and control group mean skull lengths. Because our prediction was that treatment specimens would be smaller, we utilized a single-tail alternative hypothesis, implemented in R as “greater than”.

Crown footprint area of the left lower first molar (lower M1) was represented by 25 landmarks placed around the widest portion of the crown (Figure 1C). These landmarks were used to estimate a convex hull that approximates the tooth crown area. We compared mean crown area estimates between groups using Welch’s T-Test, as described for skull length.

The decision to phenotype the lower M1 while quantifying the proteome of upper M1 was based on sampling constraints. Both molars develop simultaneously, rendering them as comparable as possible, while simultaneously accounting for the potential reciprocal destructive sampling of each method collection approach. Previous studies indicate that high power µCT has been shown to damage DNA and could possibly degrade proteins (Immel et al., 2016) and our proteomics protocol required destroying the entire upper M1. Choosing the lower molar for phenotyping also aids in future phenotypic comparisons because lower molars are usually found in higher abundance within paleontological samples, resulting in studies of phenotypic evolution and plasticity often emphasizing measures of lower molars (Bell & Jass, 2011; Burroughs, 2019; Procopio et al., 2018; Ungar, 2010). We chose an outline-based approach over traditional straight-line measurements because crown area has been shown to be a more sensitive tool for assessing size (Hopkins, 2018).

### Protein analysis by LC-MS/MS

For protein analysis we first determined if our extraction and proposed sampling worked by initially sampling only two treatment and two control specimens. After validating our protocols, we sampled the remaining two treatment and two control specimens. Proteins were isolated in 5% SDS, 100mM TEAB, 10mM DTT using a Precellys bead homogenizer for two cycles and spun at 16,000 x G for 5 minutes, which pulverized the tooth tissue. Supernatants were reduced at 55°C for 30 minutes, and cysteines were alkylated with 25mM iodoacetamide for 30 minutes at room temperature in the dark. Samples were acidified with phosphoric acid, proteins were then precipitated with 90% methanol, 50mM TEAB, and bound to S-Trap solid phase cartridges (Zougman et al., 2020). Protein precipitates were washed with 90% methanol, 50mM TEAB and digested with trypsin (20µg, Sigma-Aldrich. Trypsin, TPCK-treated; #4352157) at 47°C for two hours. We ran two blank STRAP samples per analytical sample, no protein or peptide was recovered from blank STRAP samples, except for a single keratin peptide at m/z 1082. Precipitates were then sequentially eluted with 50mM TEAB, 0.2% formic acid, and 50% acetonitrile, the 0.2% formic acid elution step was by centrifugation at 4000 x G for 1 minute each. Peptides were dried by vacuum centrifugation, desalted using HLB SPE cartridges (Waters), dried and resuspended in 20µl 0.1% formic acid/water.

Peptides were analyzed by C18 reverse phase LC-MS/MS (Thermo Nano1200) using a 2µl injection volume. HPLC C18 columns were prepared using a P-2000 CO2 laser puller (Sutter Instruments) and silica tubing (100µm ID x 15 cm) and were self-packed with 3u Reprosil resin. Peptides were separated using a flow rate of 300 nl/minute, and a gradient elution step changing from 0.1% formic acid to 40% acetonitrile (CAN) over 90 minutes, followed by a 90% CAN wash and re-equilibration steps (buffer A 0.1% formic acid/water, buffer B 90% acetonitrile/water with 0.1% formic acid). Parent peptide mass and collision-induced fragment mass information were collected using an orbital trap (Q-Exactive HF; Thermo) instrument followed by protein database searching using Proteome Discoverer 2.4 (Thermo). Electrospray ionization was achieved using spray voltage of ∼2.3 kV. Information-dependent MS and MS/MS acquisitions were made using a 50ms survey scan (m/z 375 – 1400) at 60,000 resolution, followed by ‘top 20’ consecutive second product ion scans at 15,000 resolution. Peptide and spectra false discovery rates were set to 0.01 and 0.05 FDR bins, with 10ppm MS and 0.05Da MS2 tolerances, allowing up to 2 missed tryptic cleavages, and a minimum peptide size of 6 amino acids. Variable modifications included M-oxidation, NQ-deamination, and ST dehydration. Peptide-specific label free quantitation (mapping) was performed using Proteome Discoverer 2.4 (SequestHT Percolator algorithm), linking peptides to annotated mouse proteome (Uniprot mouse 16982 sequences; 4-2019, downloaded October of 2023) and the MQ contaminant dataset for standardized protein identification. Label free abundance estimations were performed (Sequest HT LFQ). Raw data files are available on the MassIVE database using identifier MSV000098600.

### Filtering

To assess if dietary changes resulted in significantly altered protein expression, we first filtered our mapped proteomic data. Filtering was done in ProteoRE (Mehta et al., 2023). Because spectral ionization can vary between LC-MS/MS analyses (e.g., (Muntel et al., 2015; Z. Wang et al., 2024)) we initially treated our dataset as two distinct datasets, representing the initial sampling (Group 1) and additional sampling (Group 2) (Supplemental Materials). There was no expectation that Groups 1 and 2 were substantially different from one another based on the experimental procedure: groups differed only by the date of LC-MS/MS analysis. We chose a conservative approach of filtering each dataset individually and concatenating the resulting filtered datasets, ensuring that only proteins represented in both datasets were used for downstream analyses and interpretations. Each sample group was initially filtered by excluding all mapped proteins that had any one of the following criteria: A minimum false discovery rate (q-value) greater than 0.05, representation by fewer than two peptides, or three or fewer peptide spectral matches (PSMs). These criteria were selected based on proteomics field standard practices (Al-Amrani et al., 2021).

### Mapped Proteomics Analysis

To concatenate our results, a Venn analysis was performed to find the set of proteins identified as unambiguously present in both groups, resulting in a combined group (CG) dataset. This dataset reflected the shared mapped proteins between all eight (four control, four treatment) specimens in our sample. Groups 1 and 2 were further filtered to identify proteins with significant changes in protein expression between treatment and control specimens. First, proteins where at least one treatment specimen in each group had an abundance ratio adjusted p-value ≤0.05 were retained (n=135). Then, proteins that had missing data for two or more specimen comparisons were discarded (n=15). Those two datasets were subsequently concatenated via a Venn analysis to form a combined group significant (CGSig) dataset. The result is that every protein in CGSig is significantly differentially expressed in at least one treatment/control comparison in Group 1 and at least one treatment/control comparison in Group 2.

Here, the abundance ratio is a normalized ratio of estimated protein abundance, based on peak intensity, for one proteomics sample over another proteomics sample, in our case a treatment specimen over a control specimen. Because protein abundances can vary significantly in terms of magnitude between samples in label free quantification, the estimated protein abundances are commonly transformed on a Log2 scale to normalize (adjust) them (Liu & Zhang, 2021). To calculate the p-value for adjusted abundance ratios p-values we used a two-tailed Students’ T-Test. One result of this transformation is that some calculated abundance ratios will effectively become –∞ or ∞ but are represented in the dataset as values of –3.32 or 3.32. We calculated the average Log2 fold change reported in Table 1 by taking the mean of the estimated abundance ratios from samples that had significant p-values, excluding significant samples that were equal to –3.32 or 3.32. For some samples, calculated means were close to 0 and/or removal of one outlier could change the sign (and thus interpretation) of the mean. In those cases, we do not provide an average value, instead we provided the number of specimens and their direction of fold change (Up/Down) (Table 1).

**Table 1:** List of proteins with significant differential expression between low dietary protein (10%) treatment and normal dietary protein (20%) control groups. Average Log2 fold change calculated by averaging of specimens with significant change of abundance ratio (SI 1). Up=Significant specimens have positive Log2 fold change meaning increased expression of that protein. Down = Specimens have negative Log2 fold change meaning decreased expression of that protein. Proteins with blank averages are where a consistent signal was not identified due to individual variation. The number of specimens which could be individually interpreted as Up (U) or Down (D) is provided. Data are from SI 1.

| Uniprot Accession | Protein and Gene Name | GO: Process | Avg. Log2 | Up/ Down |
| --- | --- | --- | --- | --- |
| P63277 | Amelogenin, X isoform GN=Amelx | Odontogenesis | -1.52 | Down |
| O55188 | Dentin matrix acidic phosphoprotein 1 GN=Dmp1 | Odontogenesis |  | 3U/2D |
| P97399 | Dentin sialophosphoprotein GN=Dspp | Odontogenesis | 1.77 | Up |
| P43406 | Integrin alpha-V GN=Itgav | Osteogenesis | -2.66 | Down |
| Q05117 | Tartrate-resistant acid phosphatase type 5 GN=Acp5 | Osteogenesis | -1.71 | Down |
| P28481 | Collagen alpha-1(II) chain GN=Col2a1 | Osteogenesis |  | 1U/2D |
| O54974 | Galectin-7 GN=Lgals7 | Osteogenesis |  | 3U/2D |
| P27005 | Protein S100-A8 GN=S100a8 | Inflammation and Immune Response | -1.27 | Down |
| O88947 | Coagulation factor X GN=F10 | Inflammation and Immune Response | -0.23 | Down |
| Q9EPB4 | Apoptosis-associated speck-like protein containing a CARD GN=Pycard | Inflammation and Immune Response |  | 1U/1D |
| Q9JM83 | Calmodulin-4 GN=Calm4 | Inflammation and Immune Response |  | 2U/1D |
| Q9Z126 | Platelet factor 4 GN=Pf4 | Inflammation and Immune Response | 0.92 | Up |
| P16110 | Galectin-3 GN=Lgals3 | Inflammation and Immune Response | 1.10 | Up |
| P07743 | BPI fold-containing family A member 2 GN=Bpifa2 | Inflammation and Immune Response | 1.50 | Up |
| Q07456 | Protein AMBP GN=Ambp | Inflammation and Immune Response | 2.30 | Up |
| Q61176 | Arginase-1 GN=Arg1 | Inflammation and Immune Response | 2.40 | Up |
| Q9JJZ2 | Tubulin alpha-8 chain GN=Tuba8 | Cytoskeleton Formation | -2.29 | Down |
| Q8R1Q8 | Cytoplasmic dynein 1 light intermediate chain 1 GN=Dync1li1 | Cytoskeleton Formation | -1.59 | Down |
| Q9CRB9 | MICOS complex subunit Mic19 GN=Chchd3 | Cytoskeleton Formation | -1.17 | Down |
| Q91ZU6 | Dystonin GN=Dst | Cytoskeleton Formation | -0.56 | Down |
| P08032 | Spectrin alpha chain, erythrocytic 1 GN=Spta1 | Cytoskeleton Formation |  | 1U/1D |
| Q61554 | Fibrillin-1 GN=Fbn1 | Cytoskeleton Formation | 0.09 | Up |
| P15508 | Spectrin beta chain, erythrocytic GN=Sptb | Cytoskeleton Formation |  | 1U/1D |
| P55292 | Desmocollin-2 GN=Dsc2 | Cytoskeleton Formation |  | 2U/1D |
| P48193 | Protein 4.1 GN=Epb41 | Cytoskeleton Formation |  | 2U/1D |
| P10637 | Microtubule-associated protein tau GN=Mapt | Cytoskeleton Formation | 1.07 | Up |
| O35902 | Desmoglein-3 GN=Dsg3 | Cytoskeleton Formation | 1.20 | Up |
| Q7TSF1 | Desmoglein-1-beta GN=Dsg1b | Cytoskeleton Formation | 1.22 | Up |
| P97350 | Plakophilin-1 GN=Pkp1 | Cytoskeleton Formation | 1.58 | Up |
| Q02257 | Junction plakoglobin GN=Jup | Cytoskeleton Formation | 1.65 | Up |
| E9Q557 | Desmoplakin GN=Dsp | Cytoskeleton Formation | 1.80 | Up |
| G5E8K5 | Ankyrin-3 GN=Ank3 | Cytoskeleton Formation | 1.87 | Up |
| Q3UV17 | Keratin, type II cytoskeletal 2 oral GN=Krt76 | Development of Cornified Envelope |  | 3U/2D |
| A1L317 | Keratin, type I cytoskeletal 24 GN=Krt24 | Development of Cornified Envelope | 0.76 | Up |
| P04104 | Keratin, type II cytoskeletal 1 GN=Krt1 | Development of Cornified Envelope | 1.26 | Up |
| Q61781 | Keratin, type I cytoskeletal 14 GN=Krt14 | Development of Cornified Envelope | 1.46 | Up |
| P97347 | Repetin GN=Rptn | Development of Cornified Envelope | 1.47 | Up |
| Q61414 | Keratin, type I cytoskeletal 15 GN=Krt15 | Development of Cornified Envelope | 1.51 | Up |
| Q09PK2 | Retroviral-like aspartic protease 1 GN=Asprv1 | Development of Cornified Envelope | 1.70 | Up |
| Q62266 | Cornifin-A GN=Sprr1a | Development of Cornified Envelope | 1.75 | Up |
| A6BLY7 | Keratin, type I cytoskeletal 28 GN=Krt28 | Development of Cornified Envelope | 1.89 | Up |
| Q62267 | Cornifin-B GN=Sprr1b | Development of Cornified Envelope | 2.93 | Up |
| A2AQP0 | Myosin-7B GN=Myh7b | Muscle Contraction | -2.37 | Down |
| Q9QZ47 | Troponin T, fast skeletal muscle GN=Tnnt3 | Muscle Contraction | -2.34 | Down |
| **Q5SX39** | Myosin-4 GN=Myh4 | Muscle Contraction | -2.25 | Down |
| **P13542** | Myosin-8 GN=Myh8 | Muscle Contraction | -2.24 | Down |
| **Q5SX40** | Myosin-1 GN=Myh1 | Muscle Contraction | -1.91 | Down |
| **P05977** | Myosin light chain 1/3, skeletal muscle isoform GN=Myl1 | Muscle Contraction | -1.88 | Down |
| **P68134** | Actin, alpha skeletal muscle GN=Acta1 | Muscle Contraction | -1.86 | Down |
| **Q3SX28** | TREMBL:Q3SX28;Q5KR48 (Bos taurus) Tropomyosin 2 | Muscle Contraction | -1.61 | Down |
| **P13412** | Troponin I, fast skeletal muscle GN=Tnni2 | Muscle Contraction | -1.59 | Down |
| **P97457** | Myosin regulatory light chain 2, skeletal muscle isoform GN=Mylpf | Muscle Contraction | -1.42 | Down |
| **P97447** | Four and a half LIM domains protein 1 GN=Fhl1 | Muscle Contraction | -1.06 | Down |
| **A2ASS6** | Titin GN=Ttn | Muscle Contraction | -0.80 | Down |
| **Q9JI91** | Alpha-actinin-2 GN=Actn2 | Muscle Contraction | -0.48 | Down |
| **P07310** | Creatine kinase M-type GN=Ckm | Phosphorylation | -2.03 | Down |
| **P28650** | Adenylosuccinate synthetase isozyme 1 GN=Adssl1 | Phosphorylation | -1.24 | Down |
| **Q9R0Y5** | Adenylate kinase isoenzyme 1 GN=Ak1 | Phosphorylation | -0.71 | Down |
| **P29699** | Alpha-2-HS-glycoprotein GN=Ahsg | Phosphorylation |  | 2U/1D |
| **Q9R0H0** | Peroxisomal acyl-coenzyme A oxidase 1 GN=Acox1 | Regulation | -2.12 | Down |
| **P63330** | Serine/threonine-protein phosphatase 2A GN=Ppp2ca | Regulation | -1.78 | Down |
| **Q9EST5** | Acidic leucine-rich nuclear phosphoprotein 32 B GN=Anp32b | Regulation | -1.18 | Down |
| **P07759** | Serine protease inhibitor A3K GN=Serpina3k | Regulation | -1.17 | Down |
| **P28665** | Murinoglobulin-1 GN=Mug1 | Regulation | -0.88 | Down |
| **Q9Z2N8** | Actin-like protein 6A GN=Actl6a | Regulation | -0.62 | Down |
| **O55042** | Alpha-synuclein GN=Snca | Regulation | 0.29 | Up |
| **Q5FWK3** | Rho GTPase-activating protein 1 GN=Arhgap1 | Regulation |  | 1U/1D |
| **Q9D5V5** | Cullin-5 GN=Cul5 | Regulation | 0.38 | Down |
| **Q6P253** | Dermokine GN=Dmkn | Regulation | 0.89 | Up |
| **Q08189** | Protein-glutamine gamma-glutamyltransferase E GN=Tgm3 | Regulation | 1.04 | Up |
| **Q60604** | Adseverin GN=Scin | Regulation | 1.32 | Up |
| **Q9D2Q8** | Protein S100-A14 GN=S100a14 | Regulation | 1.45 | Up |
| **Q8CGR5** | Kallikrein-14 GN=Klk14 | Regulation | 1.55 | Up |
| **P21570** | Angiogenin GN=Ang | Regulation | 2.42 | Up |
| **Q62203** | Splicing factor 3A subunit 2 GN=Sf3a2 | Transcription and Repair | -0.21 | Down |
| **Q9R0Q6** | Actin-related protein 2/3 complex subunit 1A GN=Arpc1a | Transcription and Repair | 2.09 | Up |
| **P32848** | Parvalbumin alpha GN=Pvalb | Binding | -2.24 | Down |
| **P62631** | Elongation factor 1-alpha 2 GN=Eef1a2 | Binding | -2.01 | Down |
| **O35744** | Chitinase-like protein 3 GN=Chil3 | Binding | -1.82 | Down |
| **P31786** | Acyl-CoA-binding protein GN=Dbi | Binding | -1.72 | Down |
| **P81117** | Nucleobindin-2 GN=Nucb2 | Binding | -1.09 | Down |
| **O70456** | 14-3-3 protein sigma GN=Sfn | Binding | 1.09 | Up |
| **P63024** | Vesicle-associated membrane protein 3 GN=Vamp3 | Transport | -3.07 | Down |
| **Q9JLM8** | Serine/threonine-protein kinase DCLK1 GN=Dclk1 | Transport | -2.40 | Down |
| **P18872** | Guanine nucleotide-binding protein G(o) subunit alpha GN=Gnao1 | Transport | -2.35 | Down |
| **Q8VEH3** | ADP-ribosylation factor-like protein 8A GN=Arl8a | Transport | -2.18 | Down |
| **P61028** | Ras-related protein Rab-8B GN=Rab8b | Transport | -2.07 | Down |
| **P04247** | Myoglobin GN=Mb | Transport | -1.71 | Down |
| **P02089** | Hemoglobin subunit beta-2 GN=Hbb-b2 | Transport | -1.51 | Down |
| **P04117** | Fatty acid-binding protein, adipocyte GN=Fabp4 | Transport | -1.16 | Down |
| **P02088** | Hemoglobin subunit beta-1 GN=Hbb-b1 | Transport |  | 1U/1D |
| **P04919** | Band 3 anion transport protein GN=Slc4a1 | Transport |  | 1U/1D |
| **P97449** | Aminopeptidase N GN=Anpep | Transport |  | 1U/1D |
| **P14602** | Heat shock protein beta-1 GN=Hspb1 | Transport | 1.44 | Up |
| **P24668** | Cation-dependent mannose-6-phosphate receptor GN=M6pr | Transport | 2.37 | Up |
| **Q8VHI3** | GDP-fucose protein O-fucosyltransferase 2 GN=Pofut2 | Metabolism | -2.35 | Down |
| **Q5SWU9** | Acetyl-CoA carboxylase 1 GN=Acaca | Metabolism | -2.03 | Down |
| **Q9D2R0** | Acetoacetyl-CoA synthetase GN=Aacs | Metabolism | -1.93 | Down |
| **O70503** | Very-long-chain 3-oxoacyl-CoA reductase GN=Hsd17b12 | Metabolism | -1.87 | Down |
| **P19096** | Fatty acid synthase GN=Fasn | Metabolism | -1.83 | Down |
| **P21550** | Beta-enolase GN=Eno3 | Metabolism | -1.77 | Down |
| **O70250** | Phosphoglycerate mutase 2 GN=Pgam2 | Metabolism | -1.75 | Down |
| **Q8R429** | Sarcoplasmic/endoplasmic reticulum calcium ATPase 1 GN=Atp2a1 | Metabolism | -1.73 | Down |
| **Q9EQ06** | Estradiol 17-beta-dehydrogenase 11 GN=Hsd17b11 | Metabolism | -1.38 | Down |
| **P13634** | Carbonic anhydrase 1 GN=Ca1 | Metabolism | -0.92 | Down |
| **Q9WUB3** | Glycogen phosphorylase, muscle form GN=Pygm | Metabolism | -0.87 | Down |
| **Q9CQT1** | Methylthioribose-1-phosphate isomerase GN=Mri1 | Metabolism | -0.25 | Down |
| **P00920** | Carbonic anhydrase 2 GN=Ca2 | Metabolism |  | 1U/1D |
| **O89020** | Afamin GN=Afm | Metabolism |  | 1U/1D |
| **O09131** | Glutathione S-transferase omega-1 GN=Gsto1 | Metabolism | 1.22 | Up |
| **P97370** | Sodium/potassium-transporting ATPase subunit beta-3 GN=Atp1b3 | Metabolism | 1.22 | Up |
| **P47739** | Aldehyde dehydrogenase, dimeric ZDP-preferring GN=Aldh3a1 | Metabolism | 1.48 | Up |
| **Q9QYJ0** | DnaJ homolog subfamily A member 2 GN=Dnaja2 | Metabolism | 2.00 | Up |
| **Q9R099** | Transducin beta-like protein 2 GN=Tbl2 | Unknown | -1.52 | Down |
| **Q00898** | Alpha-1-antitrypsin 1-5 GN=Serpina1e | Unknown | -1.25 | Down |
| **Q9D7B7** | Probable glutathione peroxidase 8 GN=Gpx8 | Unknown |  | 1U/2D |
| **P00687** | Alpha-amylase 1 GN=Amy1 | Unknown |  | 3U/2D |
| **Q8CIT9** | Suprabasin GN=Sbsn | Unknown | 1.28 | Up |
| **Q9Z331** | Keratin, type II cytoskeletal 6B GN=Krt6b | Unknown | 1.30 | Up |
| **Q8CF02** | Protein FAM25C GN=Fam25c | Unknown | 1.81 | Up |

To identify protein functions, associated interactions, and general biological profiles represented in CG and CGSig we performed Gene Ontology (GO) enrichment analysis via the ClusterProfiler tool of ProteoRE and Pathway Enrichment Analysis via REACTOME (Croft et al., 2011; Wu et al., 2021). For GO enrichment analyses, we queried at two ontology levels, using cutoffs for p-value of 0.05 and q-value of 0.05. Outputs for GO analyses were used to assign broad categorical function to proteins (Table 1) based on Metabolic Functions, Cellular Component, and Biological Processes categories (Figure 3).

For pathway enrichment analysis via REACTOME we queried the *Mus musculus* REACTOME for the proteins found in the CGSig dataset. To calculate enrichment, the number of entities (in our case proteins) identified as belonging within a specific pathway are identified. Then the total number of entities (proteins) that could be contained in that pathway is calculated and divided by the total number of entities (proteins) from the organism (in our case *Mus*). The resulting ‘Entity Ratio’ is used to correct for pathway size to determine which pathways are overrepresented compared to a random distribution. A pathway is considered ‘enriched’ when the input from the dataset (x) is a higher proportion of the total number of entities (n) than would be expected by chance. The resulting probability estimate is represented by a p-value calculated on a 95% confidence interval and pathways with Entities p-value ≤ 0.05 are significantly enriched. Enriched pathways were then ranked based on the Entity Ratio, which represents the percentage of all *Mus* proteins in that pathway with a significant enrichment in our analysis (i.e., an Entity Ratio of 0.04 represents 4% of all *Mus* proteins known in that pathway). Pathways with higher entity ratios are ranked higher within Table 2.

**Table 2:**
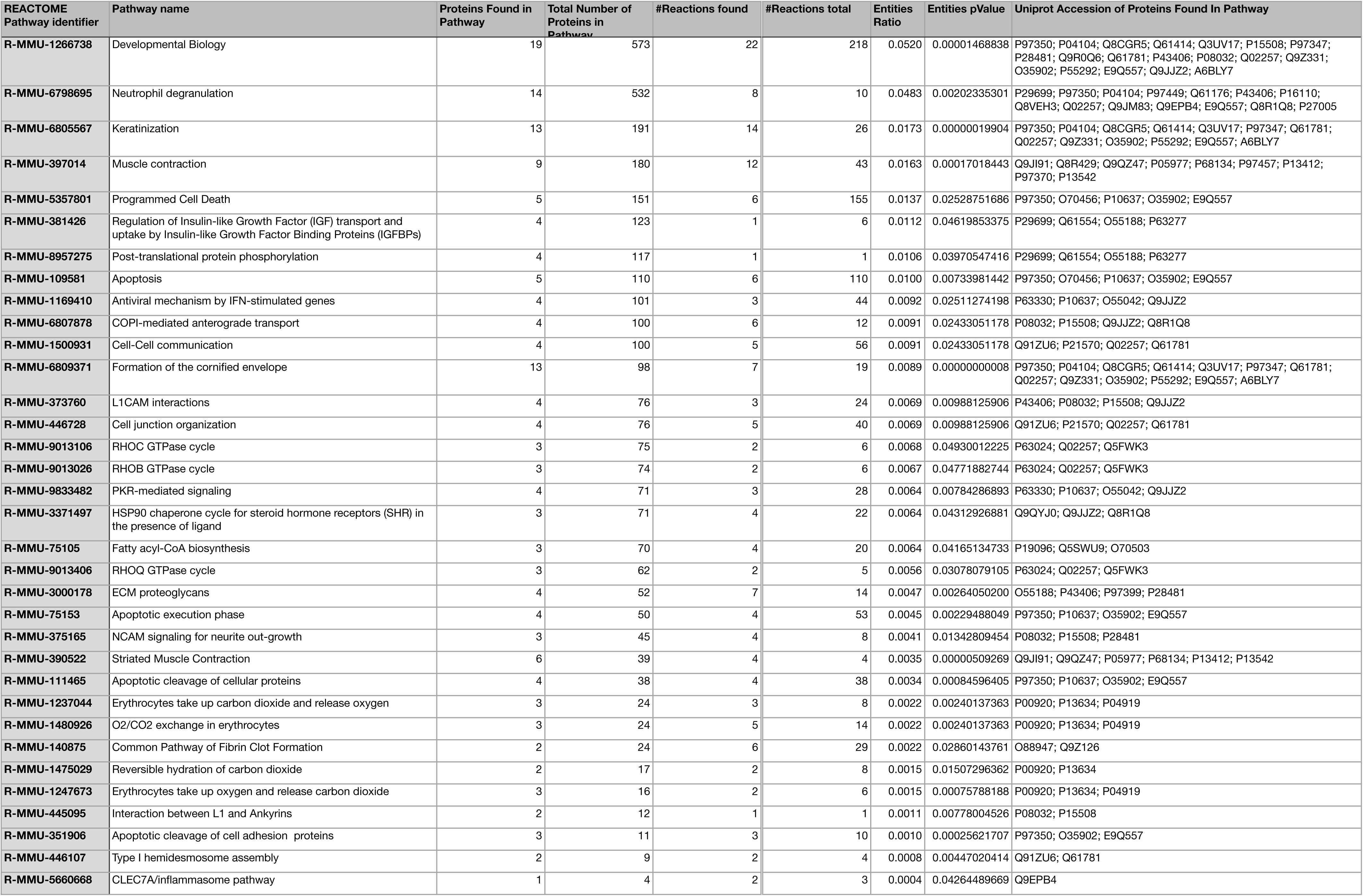
Significantly enriched REACTOME pathways derived from CGSig dataset (120 proteins). Table is ordered from highest Entities Ratio to lowest. Entities Ratio is the number of proteins from Pathway Name/Total Number of Proteins for *Mus musculus* in REACTOME database. Entities pValue is hypothesis test of chances of the number of proteins found being within pathway compared to random significant pValue = enriched pathway. Accession numbers of proteins from CGSig which belong to a pathway, not an inclusive list of all proteins known within the pathway.

| REACTOME Pathway identifier | Pathway name | Proteins Found in Pathway | Total Number of Proteins in Pathway | #Reactions found | #Reactions total | Entities Ratio | Entities pValue | Uniprot Accession of Proteins Found In Pathway |
| --- | --- | --- | --- | --- | --- | --- | --- | --- |
| R-MMU-1266738 | Developmental Biology | 19 | 573 | 22 | 218 | 0.0520 | 0.00001468838 | P97350; P04104; Q8CGR5; Q61414; Q3UV17; P15508; P97347; P28481; Q9R0Q6; Q61781; P43406; P08032; Q02257; Q9Z331; O35902; P55292; E9Q557; Q9JJZ2; A6BLY7 |
| R-MMU-6798695 | Neutrophil degranulation | 14 | 532 | 8 | 10 | 0.0483 | 0.00202335301 | P29699; P97350; P04104; P97449; Q61176; P43406; P16110; Q8VEH3; Q02257; Q9JM83; Q9EPB4; E9Q557; Q8R1Q8; P27005 |
| R-MMU-6805567 | Keratinization | 13 | 191 | 14 | 26 | 0.0173 | 0.00000019904 | P97350; P04104; Q8CGR5; Q61414; Q3UV17; P97347; Q61781; Q02257; Q9Z331; O35902; P55292; E9Q557; A6BLY7 |
| R-MMU-397014 | Muscle contraction | 9 | 180 | 12 | 43 | 0.0163 | 0.00017018443 | Q9JI91; Q8R429; Q9QZ47; P05977; P68134; P97457; P13412; P97370; P13542 |
| R-MMU-5357801 | Programmed Cell Death | 5 | 151 | 6 | 155 | 0.0137 | 0.02528751686 | P97350; O70456; P10637; O35902; E9Q557 |
| R-MMU-381426 | Regulation of Insulin-like Growth Factor (IGF) transport and uptake by Insulin-like Growth Factor Binding Proteins (IGFBPs) | 4 | 123 | 1 | 6 | 0.0112 | 0.04619853375 | P29699; Q61554; O55188; P63277 |
| R-MMU-8957275 | Post-translational protein phosphorylation | 4 | 117 | 1 | 1 | 0.0106 | 0.03970547416 | P29699; Q61554; O55188; P63277 |
| R-MMU-109581 | Apoptosis | 5 | 110 | 6 | 110 | 0.0100 | 0.00733981442 | P97350; O70456; P10637; O35902; E9Q557 |
| R-MMU-1169410 | Antiviral mechanism by IFN-stimulated genes | 4 | 101 | 3 | 44 | 0.0092 | 0.02511274198 | P63330; P10637; O55042; Q9JJZ2 |
| R-MMU-6807878 | COPI-mediated anterograde transport | 4 | 100 | 6 | 12 | 0.0091 | 0.02433051178 | P08032; P15508; Q9JJZ2; Q8R1Q8 |
| R-MMU-1500931 | Cell-Cell communication | 4 | 100 | 5 | 56 | 0.0091 | 0.02433051178 | Q91ZU6; P21570; Q02257; Q61781 |
| R-MMU-6809371 | Formation of the cornified envelope | 13 | 98 | 7 | 19 | 0.0089 | 0.00000000008 | P97350; P04104; Q8CGR5; Q61414; Q3UV17; P97347; Q61781; Q02257; Q9Z331; O35902; P55292; E9Q557; A6BLY7 |
| R-MMU-373760 | L1CAM interactions | 4 | 76 | 3 | 24 | 0.0069 | 0.00988125906 | P43406; P08032; P15508; Q9JJZ2 |
| R-MMU-446728 | Cell junction organization | 4 | 76 | 5 | 40 | 0.0069 | 0.00988125906 | Q91ZU6; P21570; Q02257; Q61781 |
| R-MMU-9013106 | RHOC GTPase cycle | 3 | 75 | 2 | 6 | 0.0068 | 0.04930012225 | P63024; Q02257; Q5FWK3 |
| R-MMU-9013026 | RHOB GTPase cycle | 3 | 74 | 2 | 6 | 0.0067 | 0.04771882744 | P63024; Q02257; Q5FWK3 |
| R-MMU-9833482 | PKR-mediated signaling | 4 | 71 | 3 | 28 | 0.0064 | 0.00784286893 | P63330; P10637; O55042; Q9JJZ2 |
| R-MMU-3371497 | HSP90 chaperone cycle for steroid hormone receptors (SHR) in the presence of ligand | 3 | 71 | 4 | 22 | 0.0064 | 0.04312926881 | Q9QYJ0; Q9JJZ2; Q8R1Q8 |
| R-MMU-75105 | Fatty acyl-CoA biosynthesis | 3 | 70 | 4 | 20 | 0.0064 | 0.04165134733 | P19096; Q5SWU9; O70503 |
| R-MMU-9013406 | RHOQ GTPase cycle | 3 | 62 | 2 | 5 | 0.0056 | 0.03078079105 | P63024; Q02257; Q5FWK3 |
| R-MMU-3000178 | ECM proteoglycans | 4 | 52 | 7 | 14 | 0.0047 | 0.00264050200 | O55188; P43406; P97399; P28481 |
| R-MMU-75153 | Apoptotic execution phase | 4 | 50 | 4 | 53 | 0.0045 | 0.00229488049 | P97350; P10637; O35902; E9Q557 |
| R-MMU-375165 | NCAM signaling for neurite out-growth | 3 | 45 | 4 | 8 | 0.0041 | 0.01342809454 | P08032; P15508; P28481 |
| R-MMU-390522 | Striated Muscle Contraction | 6 | 39 | 4 | 4 | 0.0035 | 0.00000509269 | Q9JI91; Q9QZ47; P05977; P68134; P13412; P13542 |
| R-MMU-111465 | Apoptotic cleavage of cellular proteins | 4 | 38 | 4 | 38 | 0.0034 | 0.00084596405 | P97350; P10637; O35902; E9Q557 |
| R-MMU-1237044 | Erythrocytes take up carbon dioxide and release oxygen | 3 | 24 | 3 | 8 | 0.0022 | 0.00240137363 | P00920; P13634; P04919 |
| R-MMU-1480926 | O2/CO2 exchange in erythrocytes | 3 | 24 | 5 | 14 | 0.0022 | 0.00240137363 | P00920; P13634; P04919 |
| R-MMU-140875 | Common Pathway of Fibrin Clot Formation | 2 | 24 | 6 | 29 | 0.0022 | 0.02860143761 | O88947; Q9Z126 |
| R-MMU-1475029 | Reversible hydration of carbon dioxide | 2 | 17 | 2 | 8 | 0.0015 | 0.01507296362 | P00920; P13634 |
| R-MMU-1247673 | Erythrocytes take up oxygen and release carbon dioxide | 3 | 16 | 2 | 6 | 0.0015 | 0.00075788188 | P00920; P13634; P04919 |
| R-MMU-445095 | Interaction between L1 and Ankyrins | 2 | 12 | 1 | 1 | 0.0011 | 0.00778004526 | P08032; P15508 |
| R-MMU-351906 | Apoptotic cleavage of cell adhesion proteins | 3 | 11 | 3 | 10 | 0.0010 | 0.00025621707 | P97350; O35902; E9Q557 |
| R-MMU-446107 | Type I hemidesmosome assembly | 2 | 9 | 2 | 4 | 0.0008 | 0.00447020414 | Q91ZU6; Q61781 |
| R-MMU-5660668 | CLEC7A/inflammasome pathway | 1 | 4 | 2 | 3 | 0.0004 | 0.04264489669 | Q9EPB4 |

### Preliminary Test of Developmental Archive

Prior research and the design of our study allowed us to conduct a preliminary investigation of whether enamel and dentin proteomes represent an archive of protein expression during mineralization, rather than proteins expressed earlier in tooth development or at the time of specimen collection. We queried the Mouse Gene eXpression Database (GXD) to determine if a subset of proteins was expressed at developmental stages prior to the onset of mineralization, during mineralization, or after mineralization was complete. First, we investigated proteins known to be expressed during tooth mineralization (Pandya et al., 2017). The study of Pandya et al. (2017) reported 24 proteins present during tooth mineralization, we identified 20 of those 24 proteins in our CG dataset. We also investigated proteins related to odontogenesis/osteogenesis, immune, and actin-based myosins from our CGSig dataset. A protein’s associated gene being expressed during mineralization, but not earlier or later in time, would support the argument that tooth proteomes represent a specific window of development during amelogenesis. The table containing the proteins queried in the GXD is within Supplemental Materials.

## Results

### Phenotyping

We found no significant differences in size between treatment and control groups (Table 3). Mean skull length for the treatment group was 19.20mm and control group was 19.03mm (Figure 2A). Mean crown area for treatment group was 1.03mm^2^ and control group mean was 1.02mm^2^ (Figure 2B).

**Figure 2:**
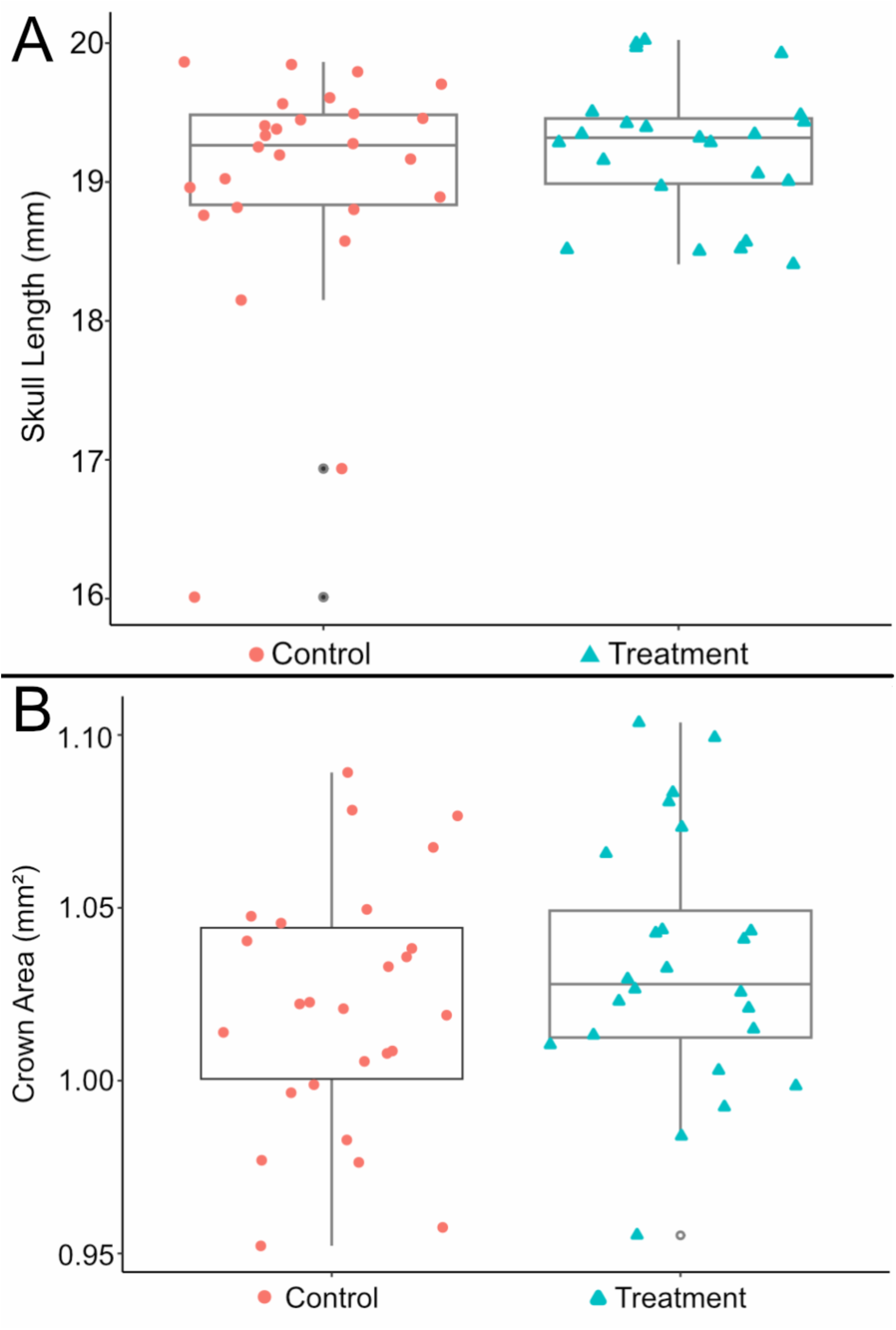
Results of phenotypic investigation. A: Estimates of straight line skull length. B: Estimates of crown area.

**Figure 3:**
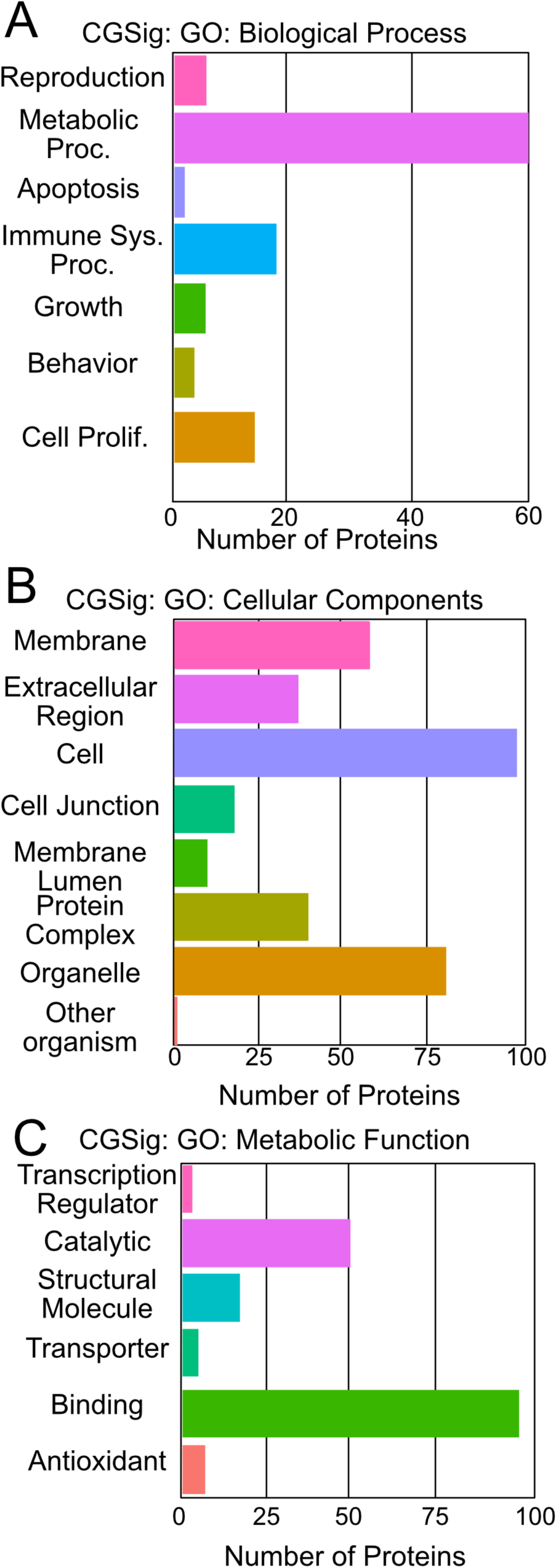
Most enriched pathways from Gene Ontology (GO) analysis of CGSig. A: Biological Processes, B: Cellular Components, C: Metabolic Function

**Table 3:** Welch’s Two Sample T-Tests to test for significant difference in mean between treatment and control group crown area and skull length.

| *Crown area by Group* |  |  |  |
| --- | --- | --- | --- |
|  | t = -1.1575 | df = 47.474 | p-value = 0.8736 |
| *95% Confidence Intervals* | -0.02914971 | Infinity |  |
| *Mean Control* | 1.021672mm<sup>2</sup> |  |  |
| *Mean Treatment* | 1.033573mm<sup>2</sup> |  |  |
| *Skull Length by Group* |  |  |  |
|  | t = -86423 | df = 41.17 | p-value = 0.8038 |
| *95% Confidence Intervals* | -0.5085681 | Infinity |  |
| *Mean Control* | 19.02755mm |  |  |
| *Mean Treatment* | 19.20011mm |  |  |

### Proteomics

Within G1, a total of 2189 unique proteins were mapped (Supplemental Materials). 1622 unique proteins were mapped for G2 (Supplemental Materials). The combined group (CG) of proteins shared between G1 and G2 was 1469 unique proteins (Supplemental Materials). Of the combined group, there were a total of 120 proteins with significant differential expression (fold change) (CGSig; Table 1; Figure 4; Supplemental Materials Figures 1-6).

**Figure 4:**
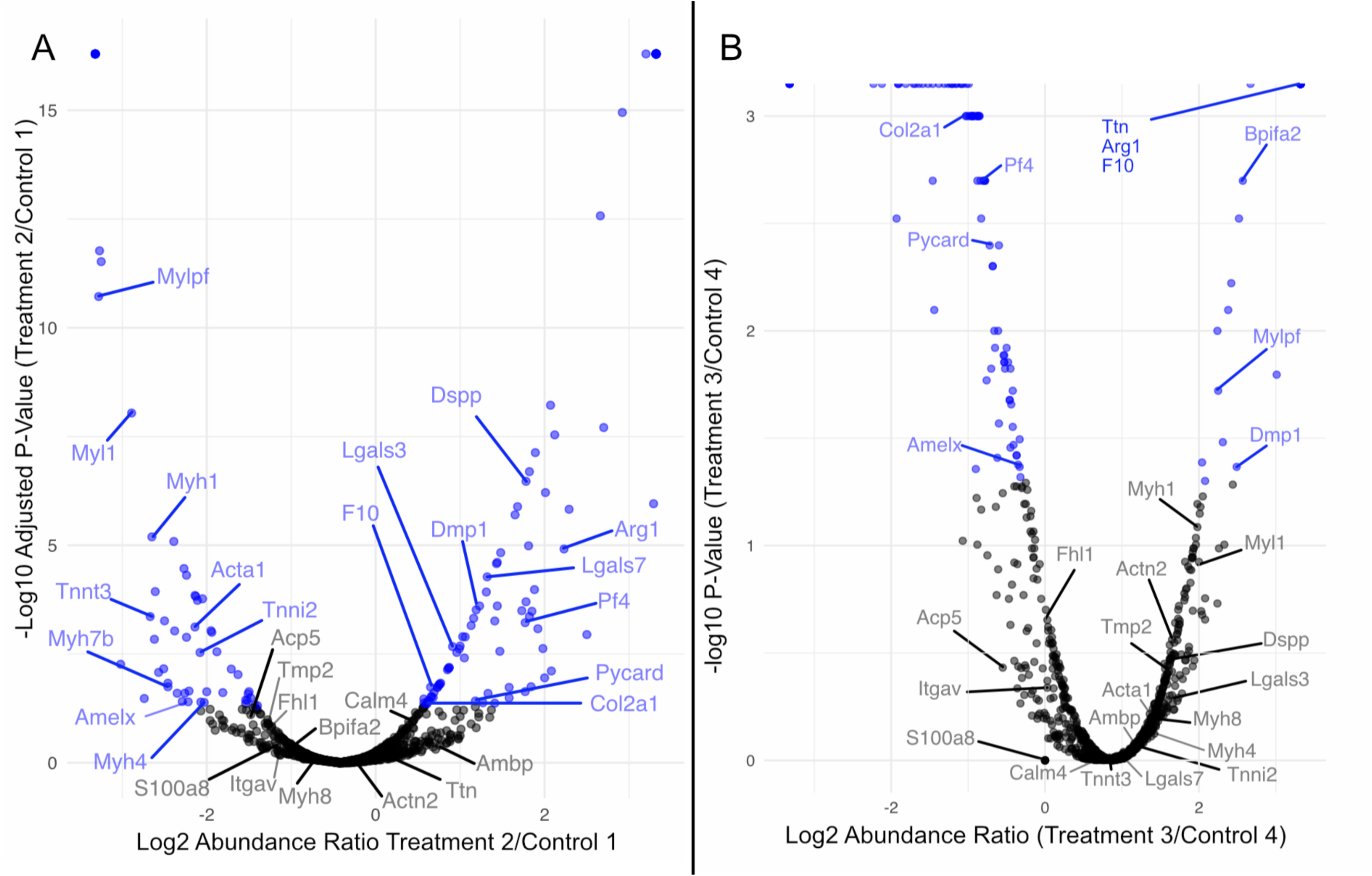
Volcano of specimen proteomics data exemplified by selected comparison from each group. A: From Group 1 Treatment 2 over Control 1. B: From Group 2 Treatment 3 over Control 4. Blue dots show Adjusted Abundance Ratio P-values ≤ 0.05. Labeled dots are proteins related to osteogenesis, odontogenesis, inflammation and immune response, and muscle contraction. Volcano plots of remaining 6 comparisons are provided in Supplemental Materials.

Gene ontology profiling revealed that both CG and CGSig are comprised of sets of proteins primarily associated with the Binding (in Molecular Function), the Cell generally (as opposed to a specific cellular component), and the Biological Process of Metabolism (Figure 3; Supplemental Materials). Pathway enrichment analysis via REACTOME identified 387 biological pathways associated with significantly differentially expressed proteins (Supplemental Materials). Of those 387, 34 were significantly enriched (p ≤ 0.05) (Table 2). Approximately 47% of matched proteins are represented in the top 6 enriched pathways (Entity Ratio ≥ 0.01) (Table 2; Supplemental Materials).

### Odontogenesis and Osteogenesis Proteins

A total of 8 proteins in CG were identified as belonging to biomineralization seven proteins associated of them had significant differential expression (Log2 fold change) between treatment and control groups (Table 1; Figure 4). In low-protein treatments, the major enamel-forming protein, Amelogenin X (AMELX) had an average –1.52-fold change in expression. One of the two major dentin-forming proteins, Dentin sialophosphoprotein (DSPP) had a 1.77-fold increase in expression. The other dentin-forming protein, Dentin matrix acidic phosphoprotein 1 (DMP1) did not have a clear direction of differential expression; three treatment comparisons showed an increase in protein expression, while two had a decrease in expression. For osteogenesis, Integrin alpha-V (ITGAV) had –2.66-fold change and Tartrate-resistant acid phosphatase type 5 (ACP5) had –1.71-fold change. The remaining two osteogenic proteins Collagen alpha-1(II) chain (COL2A1) and Galectin-7 (LGALS7) had variation that made interpretation unclear. For COL2A1, one specimen had increased and two had decreased expression. For LGALS7, three were increased and two decreased.

### Inflammation and Immune Response

A total of 229 proteins were associated with Immune System Response. Nine proteins associated with inflammation and immune system response had significant differential expression (Log2 fold change) between treatment and control groups (Table 1; Figure 4). Two of the nine proteins, S100-A8 (S100A8) and Coagulation factor X (F10), had a – 1.27 and –0.23 fold change in treatment groups, respectively. Five of the nine proteins had increases in expression: 0.92 for Platelet factor 4 (PF4), 1.10 for Galectin-3 (LGALS3), 1.50 for BPI fold-containing family A member 2 (BPIFA2), 2.30 for Protein AMBP (AMBP), and 2.40 for Arginase-1 (ARG1). Two of the immune proteins, Apoptosis-associated speck-like protein containing a CARD (PYCARD) and Calmodulin-4 (CALM4), lacked consistent signal between pairwise specimen comparisons. For PYCARD, one specimen showed increased expression and one showed decreased expression. For CALM4, two specimens were increased in expression and one decreased in expression.

### Muscle Contraction

A total of 42 proteins were associated with muscle contraction 13 of those proteins had significant differential expression. All of them had decreased expression for treatment specimens (Table 1; Figure 4). Seven of the 13 proteins are actin-based myosins (Myosins 1, 4, 7B, 8, Myosin Light chain 1, Tropomyosin 2, and Myosin regulatory light chain 2), with the remaining six being actin-specific proteins (Troponin T, Actins Alpha 1 and Alpha 2, Titin, and Four and a half LIM domains protein 1).

### Preliminary Test of Developmental Archive

Genes associated with two of 20 tooth proteins previously found during mineralization (Pandya et al., 2017) are also found in tooth tissues prior to the onset of amelogenesis based on a query of GXD. Those genes are *Alpl* and *Itgb1*. Both are found only in the tooth developmental stage immediately prior to amelogenesis (TS21 associated with embryonic days 12.5-14). The genes associated with the remaining 18 proteins are found only in developmental stages associated with molar amelogenesis (TS22+, embryonic day 15 through postnatal day 8). Querying genes associated with our proteomic sample’s differentially expressed immune and actin-based myosin proteins within GXD revealed that immune genes are universally expressed within tooth tissues during all reported stages until adulthood. Actin-based myosins were present from the onset of mineralization forward. Two of the actin-based myosins (*Actn2* & *Fhl1*) were reported as definitively absent from tooth tissues at E14.5 (i.e., just prior to mineralization) (Visel et al., 2004). Additionally, one gene (*Myh1*) was present in mineralization stages but was definitively absent from postnatal week 6-8 aged mouse specimens (Freeman et al., 1998).

## Discussion

### Phenotyping

Our hypotheses were that specimens in the treatment group would have reduced skull length and M1 size, results we did not recover. Both phenotypic results were unexpected because many studies have reported variation in skull size associated with nutritional quality in rats, mice, and pigs (e.g., (Barbeito-Andrés et al., 2016; Dickerson & Hughes, 1972; Lobe et al., 2006; McCance et al., 1961; Miller & German, 1999; Pucciarelli, 1980, 1981), and four previous studies reported variation in the size of the M1 due to dietary changes (Paynter & Grainger, 1956; Holloway et al., 1961; Paynter & Grainger, 1961; Shaw & Griffiths, 1963). Why then did we not recover similar signals?

First, we note that these previous studies sampled specimens at different ontogenetic stages than we did, most sampled specimens that were older than ours (40-days postnatal to 10-months postnatal), a single study (Barbeito-Andrés et al. 2016) sampled specimens younger than ours (embryonic day 18 to postnatal day 1). Considering skull length, mice grow continuously until they reach sexual maturity and rapidly between birth and weaning. It is possible that any differences in skull length were simply not manifested at the ontogenetic stage we sampled. This does not explain, however, the lack of difference in M1 size reported previously, because M1s are determinate in growth and cease to grow by postnatal day 8 (Ungar, 2010; Pandya et al, 2017). In comparison to our study design using 10% dietary protein as the low-protein treatment, some other studies used dietary protein quantity as low as 8% (e.g., Paynter & Grainger, 1956; Holloway et al., 1961; Paynter & Grainger, 1961; Shaw & Griffiths, 1963; Miller & German 1999). This small additional reduction in dietary protein may be sufficient to trigger effects. Finally, we recognize that different dental and cranial traits are impacted by dietary protein more readily than others. Cranial traits have been reported to be less impacted by reduced dietary protein than postcranial traits (e.g., McCance et al. 1961; Lobe et al. 2006) and M1 size may be less impacted than other traits of the M1 (e.g., angle of cusps) (Paynter & Grainger, 1956; Holloway et al., 1961; Paynter & Grainger, 1961).

At present we can only conclude that there is no difference in M1 size or skull length of specimens from our groups. However, our proteomic results (discussed in detail below) inform us that development of the tooth was perturbed in ways that likely impact other aspects of tooth phenotype. For instance, changes in relative enamel or dentin thickness would not be recovered by our estimates of size, because these are internal structural changes to tooth composition. Future more comprehensive phenotyping efforts may reveal significant differences.

### Developmental Archive

Results from GXD queries support the argument that our tooth proteome dataset primarily represents gene expression during mineralization and not prior to mineralization. For example, genes for actin-based myosins of interest within our dataset are not expressed in mouse teeth prior to the onset of amelogenesis. While it is not possible to say whether the measured immune-system proteins in our proteomic sample were expressed during mineralization or at the time of euthanasia, previous studies (e.g., Green et al., 2019; Jagr et al., 2019) have identified immune and inflammation related proteins incorporated in tooth tissues, either within enamel itself or within other tissues of the enamel-dentin junction, such as salivary glands. In our case, we recover several of the immune related proteins reported by those studies, including S100A8 and CALM4, potentially supporting the idea that molar enamel represents an archive of gene and protein expression during amelogenesis. This conclusion is further supported by the fact that the gene for differentially expressed MYH1 in our dataset is not expressed after mineralization, indicating our MYH1 signal likely represents protein expression during mineralization rather than a later time point (Freeman et al. 1998). Further, we would not expect expression of amelogenesis-specific genes or proteins after the end of mineralization, because of the cessation of proliferation of ameloblasts and lack of vascularization within the fully mineralized tooth (Alghadeer et al., 2023; Nanci, 2007). While this hypothesis requires further directed experimental validation, our results support the concept that the recovered tooth proteome represents a limited window of development.

### Odontogenic Proteins

We predicted that proteins associated with enamel and dentin formation would be altered by our feeding experiment, and specifically that they would be decreased in expression. The major enamel forming protein, AMELX, was significantly reduced in expression for treatment specimens (Table 1; Figure 4). We anticipated that our dietary protein reduction would result in decreased tooth size, associated with decreased AMELX expression, because AMELX is a necessary component for the formation of enamel. While changes in AMELX met our expectations, measures dentin forming protein expression and lower M1 size did not.

In the case of DMP1, there is not a clear signal to interpret whether expression was increased or decreased in our proteomics sample. This highlights the challenges of drawing interpretations from proteomic data where individual variation can influence the overall interpretation. This challenge is recognized by the field of quantitative proteomics, but still represents an area where increased research efforts will be needed (Al-Amrani et al., 2021; Chantada-Vazquez et al., 2022; Liu & Zhang, 2021; Steward et al., 2023). A clearer interpretation of DMP1 expression, would be helpful for constructing future hypotheses. For instance, decreased expression of DMP1 should lead to decreased expression of DSPP and dentin hypomineralization, suggesting that DMP1 and DSPP expression contribute significantly to dentinogenesis imperfecta (Orsini et al., 2014; Shi et al., 2020). Being able to robustly identify such patterns or, at minimum, make supported interpretations based on the variable evidence, will enhance the utility of future quantitative proteomic studies.

Our finding of increased DSPP expression seemed initially counterintuitive. However, this result is supported by a recent study of protein expression in a hypomineralized enamel defect found in humans (Mukhtar et al., 2022). In this study of hypomineralized molars, the enamel defect impacts the first permanent molars of children and results in a significant reduction in mineral density from normal teeth (Mukhtar et al., 2022). This reduced density was associated with downregulation of AMELX, upregulation of DSPP, but no reported differences in DMP1 expression (Mukhtar et al., 2022).

The causal mechanism for the pattern of increased DSPP expression is unknown. The general role of DSPP is to control the conversion of dental pulp cells into odontoblasts via binding with Integrin beta 6 (ITGB6) (Ritchie, 2018; Wan et al., 2016). Previous work indicated that mice with either a DSPP heterozygous (DSPP^+/-^) or DSPP knockout (DSPP^-/-^) genotypes experience dentin dysplasia and dentinogenesis imperfecta due to haploinsufficiency of DSPP (Shi et al., 2020). Haploinsufficiency suggests that dentin is impacted when DSPP expression is decreased but does not indicate what phenotype results from elevated DSPP expression. Amelogenesis imperfecta enamel is typically thin and chalky while dentin appears to be normally mineralized, suggesting that DSPP overexpression does not result in dentin hypermineralization, but this has not been experimentally validated and dentin structure was not reported by Mukhtar et al. (2022).

Our proteomic results appear consistent with protein expression patterns associated with amelogenesis imperfecta and not dentinogenesis imperfecta, based on the shared expression changes for AMELX and DSPP (Mukhtar et al., 2022; Orsini et al., 2014; Shi et al., 2020). It is unlikely that both amelogenesis and dentinogenesis imperfecta are simultaneously present within a single specimen’s dentition. Only a single study has reported compounded presence of amelogenesis and dentinogenesis imperfecta, which occurred in an MSX2 knockout transgenic line (Aioub et al., 2007).

It is possible that the signal here is due to experimental design of this study, where enamel and dentin are pooled together. The concern is that dentin constitutes a larger amount of the overall tooth and could ‘swamp’ the proteomic signal. However, this appears unlikely due to consistent signals of expression changes for many proteins not related to dentin. In cases where a set of proteins ‘swamp’ the signal, it is due to exceptionally high abundance of certain proteins, e.g., raw blood is dominated by ∼20 proteins related to hemoglobin, swamping out other protein signals (Molloy et al., 2023). In our case, recovery of 1469 proteins and 120 significantly different proteins associated with numerous processes suggests that our signal is not driven by one tissue or category of proteins within the pooled tissue sample.

Thus, we conclude that our measures of differentially expressed amelogenesis-associated proteins indicate that the development of thinner and less mineralized dental enamel could be a phenotypically plastic response to reduced dietary protein during early development. Such a result would help explain why there are disparate signals between proteomic and phenotypic results: while tooth size is not affected, tooth or microstruture composition may be. To test this hypothesis, it is necessary to utilize synchrotron scanning or destructive sampling in the form of histology. For now, it remains to be tested if the mouse molars from our study have amelogenesis imperfecta.

### Immune and Inflammation Proteins

Previous work on mapping protein expression across micro-sampled enamel sections of pig molars had suggested that there was possibility of recovering immune and inflammation related proteins from mineralized dentition (Green et al., 2019). In their case, Green et al. (2019) were constructing a detailed map of proteomic expression associated with amelogenesis. Their reported immune system proteins were localized from enamel which came from along the enamel-dentin junction (Green et al., 2019). Their study was not designed nor attempted to induce differential protein expression based on experimental procedures. In our case, we were unsure if we would recover immune and inflammation proteins because we created a tissue-averaged signal by crushing and processing the entire M1. By finding these proteins in our proteomics sample and recovering differential expression of them, we present a novel result of immune response to an environmental change. Of the nine immune or inflammation response proteins with significant fold change, five were increased in expression for treatments relative to controls, two were reduced in expression, and two had mixed interpretations (Table 1).

Seven of the nine proteins are primarily associated with neutrophil degranulation, including transport and proliferation of neutrophils. Of these seven, five were increased in expression. However, Calmodulin 4 (CALM4) had a mixed interpretation and coagulation factor X (F10) was reduced in expression. Neutrophils function as critical, but specialized, immune system response cells. Neutrophils contain granules of multiple types (azurophilic, specific, ficolin-rich, tertiary, and secretory), that target specific threats and/or regulate immune system response to specific infectious threats (Eichelberger & Goldman, 2020; Othman et al., 2022; Yin & Heit, 2018).

The precise nature of what was being targeted by immune system activation is unknown. However, the result of F10 being reduced in expression may provide some insight into the potential infectious threat. Deficiency of F10 has been implicated as part of an immune response to the common, antibiotic-resistant, bacterium *Acinetobacter baumannii* (Choby et al., 2019). In those cases, F10 deficiency is indicative of an increased abundance of neutrophils and macrophages (Choby et al., 2019). Thus, though F10 is decreased in expression, relative to the increase in five other neutrophil degranulation proteins, the combination of patterns supports a conclusion that our treatment group had higher immune system response than our control group. Future research efforts to systematically compare immune-related proteomic signatures from mineralized structures to standardized health monitoring tools could prove fruitful.

We also recovered changes in PYCARD and S100A8 within our proteomics sample, that are indicative of an inflammation response (Table 1). Studies have indicated that PYCARD directly mitigates inflammation when upregulated and contributes to inflammation when downregulated (Sartoretto et al., 2019; Wittmann et al., 2021). In our sample, one treatment specimen showed a moderate increase in PYCARD expression over controls, and one showed a moderate decrease (Table 1). Importantly, deviation in either direction suggests that there is either an increased response to inflammation (increased expression) or increased inflammation present (decreased expression) (Sartoretto et al., 2019; Wittmann et al., 2021). Reduced expression is recovered for S100A8, consistent with an ongoing physiological response to inflammation (S. Wang et al., 2018). These patterns of differential expression may represent increased inflammation response in some of the treatment specimens, as compared to control specimens. Due to the limited nature of our data and potential variation in inflammation response between specimens, these results should be interpreted cautiously.

Because of our controlled experimental design, we postulate that reduced dietary protein is the cause of increased inflammation. A recent study investigating the impact of low dietary protein during gestation indicates that intrauterine inflammation can occur and result in increased inflammation present in the offspring of Syrian golden hamsters (Mohammed et al., 2023). While Mohammed et al. (2023) focused on measuring inflammation of the liver of the offspring, the connection between low dietary protein during embryonic and postnatal development and increased inflammation was strongly established. Our study and results further support this connection. Future studies should aim to systematically investigate proteomic signals of mineralized structures along with standardized inflammation panels. This would further establish the connection between dietary protein, inflammation, and the signals archived within mineralized structures.

### Muscle Contraction Proteins

All thirteen significantly modified Muscle Contraction pathway proteins were reduced in expression. They were predominantly actin-based myosins, which play a critical role in cellular movement and structure formation by acting as motor molecules (Guhathakurta et al., 2018). Actin-based myosins are broadly implicated in the proper development of many tissues, including dentition (Du et al., 2024; Guhathakurta et al., 2018; Luis & Schnorrer, 2021). During dental development actin-based myosins contribute to the proper formation of enamel rods by transporting ameloblasts (Duverger & Morasso, 2018). Lower expression of actin-based myosins, including some of those that are differentially expressed in our study (e.g., Myosin-1, Myosin-4, Myosin-8, and Tropinin 1) are associated with the syndromic form of amelogenesis imperfecta (Duverger & Morasso, 2018). To date, no proteomic study of dentition has recovered differential expression of actin-based myosins, but none of the previous dental proteomic studies attempted to experimentally induce differential expression of these proteins. Future investigations should center on understanding the distribution of actin-based myosins within the tooth and the correlation between protein expression in teeth and other tissues when dietary-protein is reduced.

### Potential Pathways for Phenotypic Plasticity

We recovered 34 significantly enriched REACTOME pathways from our 120 significantly differentially expressed proteins. Among those pathways, two of the top six most enriched pathways, regulation of insulin-like growth factor and muscle contraction, have proteins that play many roles during morphogenesis.

Insulin-like growth factor plays a role in a number of different developmental processes, including odontogenesis, bone development, organ development, and brain development (Baroncelli et al., 2017; Brown et al., 2017; Chen et al., 2009; Dodington et al., 2021; Luo et al., 2020; Montivero et al., 2021; Oyanagi et al., 2019; Vassilakos et al., 2019). Thus, regulation of IGF is a complex set of processes with many different factors involved in regulating expression in different tissues. This includes regulation by thyroid growth hormone (Yakar & Isaksson, 2016) and regulation via phosphorylation of binding proteins to aid in transport, activation, and inhibition (Chrudinova et al., 2024; Dong et al., 2022; Huttlin et al., 2010; Palma-Lara et al., 2023; Tagliabracci et al., 2015). Given the importance of IGF to multiple developmental processes, regulation of IGF is a probable candidate for a mechanism underlying phenotypic plasticity. Experimental manipulation of IGF1 and IGF2 during odontogenesis has revealed systematic changes to the size and number of cusps of developing molars (Oyanagi et al., 2019). The connection between IGF1 and IGF2 gene expression and proteins associated with IGF regulation, which includes Amelogenin X, is not currently well understood (Bansal et al., 2012; Oyanagi et al., 2019; Pandya & Diekwisch, 2021). However, regulation of insulin-like growth factor offers a promising pathway for future investigation, particularly in elucidating responses of IGF gene expression and associated regulatory factors to environmental perturbations, such as poor diet.

Similarly, our collection of proteins from the muscle contraction pathway are primarily actin-based myosins. The function of myosins as motor proteins is well documented, but the specific nature of how myosins interact during odontogenesis is poorly understood (Du et al., 2024; Guhathakurta et al., 2018). Previous proteomic studies of teeth have not reported differential expression of actin-based myosins, making this result unexpected. Simultaneously, those studies were not attempting to experimentally induce plastic responses like our study. The connection of actin-based myosins with body size and potentially with amelogenesis imperfecta suggests that future investigations may be fruitful. These future studies would hopefully confirm the results we recovered and further explore the relationship between protein expression and phenotypic plasticity.

### Conclusions

Proteomic expression results supported our prediction that halving dietary protein during embryogenesis and early postnatal development would alter expression of proteins recovered from mouse molars. Specifically, we identified 120 differentially expressed proteins associated with a reduction of dietary protein during embryonic and early postnatal development. Changes in dietary protein have been proposed to result in phenotypically plastic changes in molar size and skull length. Our study did not recover significant differences between normal and low protein groups for either a measure of molar size or skull length. The discordance between positive proteomic results and negative phenotypic results suggests that phenotypic differences may be more nuanced that initially predicted.

We recovered significant changes in proteins associated with dental development, that are primarily within the pathway associated with regulation of Insulin-Like Growth Factor. The connection between IGF and dental proteins remains to be further investigated, but changes in expression in this pathway could directly influence tooth structure. We also identified systematically reduced expression for proteins in the Muscle contraction pathway, specifically actin-based myosins, a novel discovery for tooth-derived proteomics data. Actin-based myosins are broadly implicated in vertebrate development and are correlated with dental and general body development. Taken together, it is possible that we perturbed tooth phenotypes, such as enamel or dentin thickness or density, that are not captured by our measures of craniodental size.

Fortunately, the study presented here is part of a larger research program aimed at more comprehensive phenotyping efforts. The results of this study will allow us to focus on investigating phenotypic hypotheses derived from our proteomics results (e.g., testing for amelogenesis imperfecta). As well as continuing to investigate other aspects of craniodental morphology, to include assessment of size of the second and third molars and shape of all three molars, along with cranial traits such as depth and width of the cranium. Inclusion of these more comprehensive phenotypic data may recover differences between groups that were not recovered from the data reported here.

Our controlled feeding experiment also induced increased immune system activation and inflammation response, as evidenced by increased expression of proteins in the Neutrophil degranulation pathway. While this result does not directly inform us about phenotypic plasticity, that we could potentially to derive such information from a fully mineralized teeth could prove useful for studying the biology of deceased or extinct organisms. New studies aimed at microsampling enamel, dentin, and along the enamel-dentin junction to determine both the source of these signals and which tissues contain them are necessary.

Finally, we propose that proteomic quantification of non-experimental organisms will prove fruitful and predict that dietary changes in wild settings will change enamel and dentin formation by altering aspects of the IGF regulatory pathway and potentially expression of actin-based myosins. Future research efforts then should focus on elucidating the connection between IGF gene expression and enamel and dentin protein expression; determining the role of actin-based myosins in tooth and skeletal development; and comparing the immune and inflammation signals in proteomic data to classically used tools (e.g., blood panels and cortisol screening). With a combination of phenotyping and proteomic investigations it will become clearer precisely what traits are more likely to be perturbed, and what sections of these pathways are causal to those changes. In turn, this will enhance the utility of quantitative proteomics for investigating organismal biology more broadly.

## Acknowledgments

We thank the Stony Brook Division of Laboratory Animal Research (DLAR), particularly Laurie Levine, for animal husbandry and management. We thank Dr. John Haley of the Stony Brook University Proteomics Core Facility, for sample preparation and initial post-processing of proteomics data. We thank the New York Institute of Technology Visualization Center staff, Dr. Brian Beatty and Kelsi Hurdle, for providing technical support in scanning and reconstruction of µCT scans. We thank the Stony Brook University Center for Inclusive Education staff: Lisa Ospitale, Diana Champney, and Erica Valdez, and Stony Brook IRACDA PIs, Dr. Carol Carter and Karian Wright, for financial and administrative support.

Author RWB was funded via an Institutional Research and Academic Career Development Award (IRACDA) made to Stony Brook University from the National Institute of General Medical Sciences of the National Institutes of Health [K12GM102778], and startup funding to NSV from Stony Brook University. The content is solely the responsibility of the authors and does not necessarily represent official views of the National Institutes of Health.

## Data availability statement

All mapped and analyzed proteomics data and phenotypic data analyzed, along with R-scripts and ProteoRE workflow are included as Supplemental Materials. These Supplemental Materials will be made available publicly upon publication via Dryad Digital Repository DOI: 10.5061/dryad.kh18932j0. Raw proteomics data will be made publicly available on the MassIVE database with Dataset Identifier: MSV000098600.

## Ethics approval statement

Live animal experiments were conducted in accordance with guidelines and regulations from Stony Brook University Institutional Animal Care and Use Committee (IACUC). Research was conducted in accordance with plan and procedures in approved protocol SBU IACUC 2023-0014.

## Patient consent statement

Not Applicable.

## Permission to reproduce material from other sources

Not Applicable.

## Clinical trial registration

Not Applicable.

## Supplemental Figures

Volcano Plots from Burroughs et al. (In Review) Tooth Proteomics

Low-dietary protein (10%) sample specimen over control (20% protein) sample specimen. Blue dots show adjusted p-value is ≤ 0.05. Labeled proteins refer to Table 1 in Burroughs et al. (In Review).

**1:**
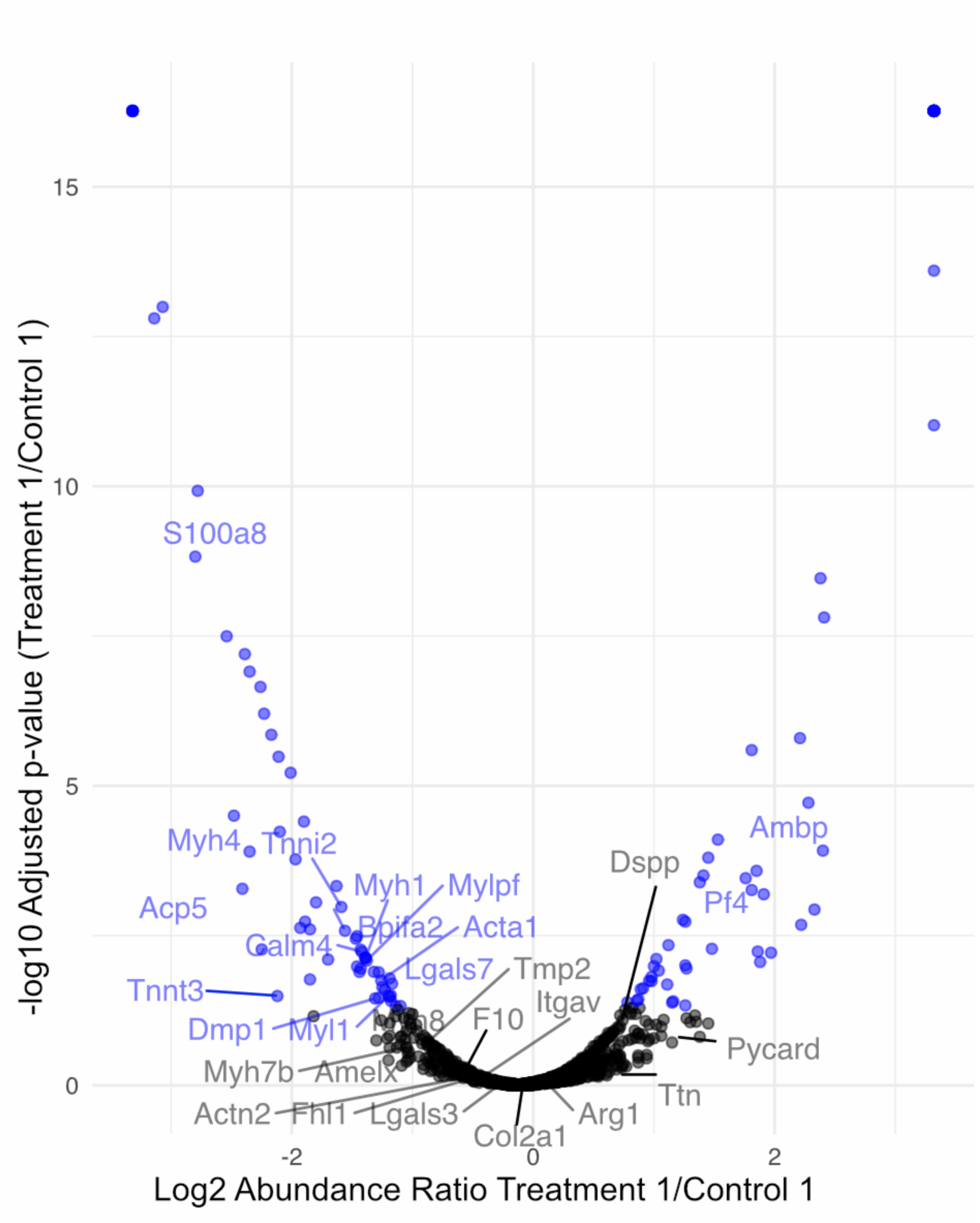
Treatment 1 Over Control 1.

**2:**
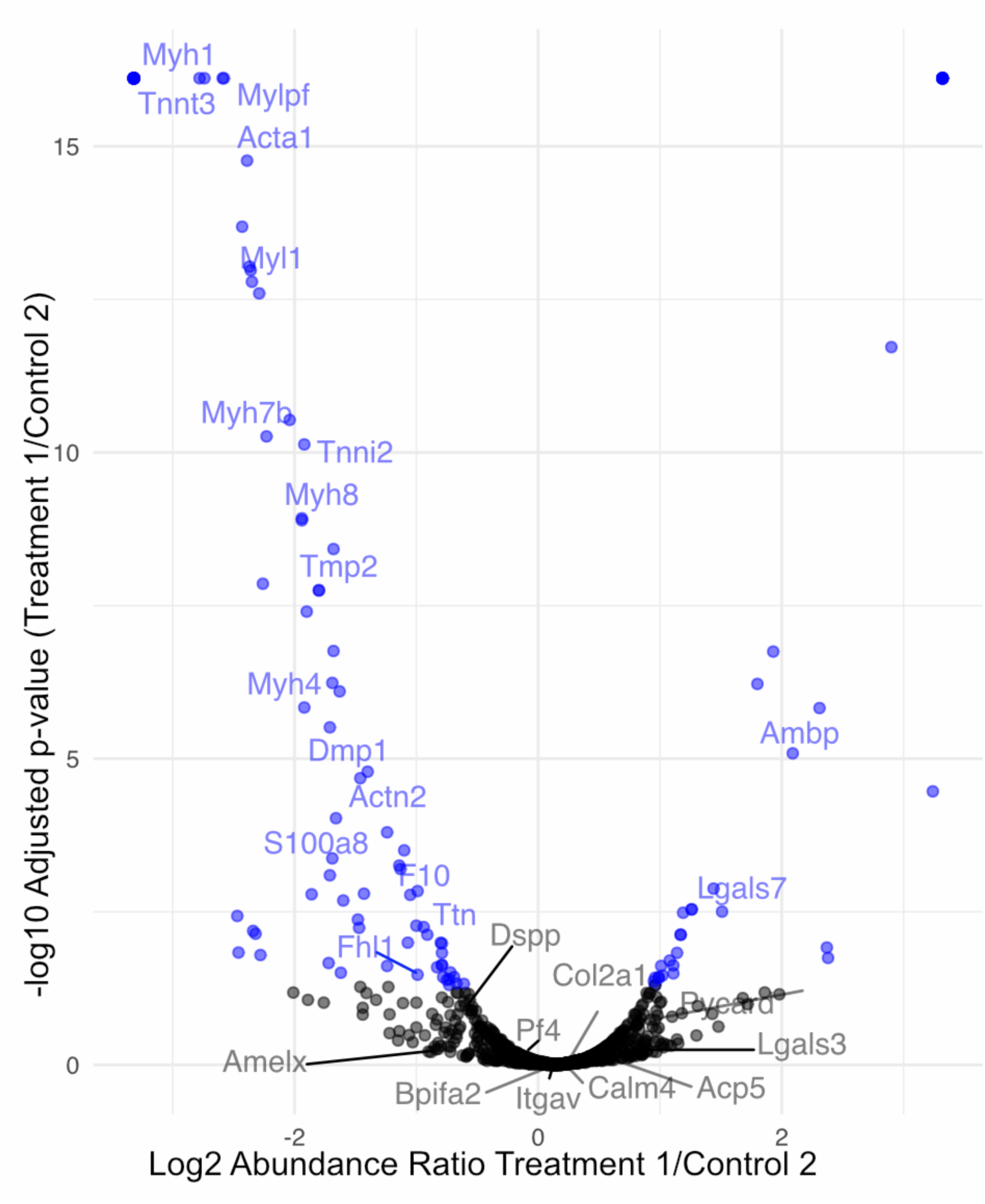
Treatment 1 Over Control 2.

**3:**
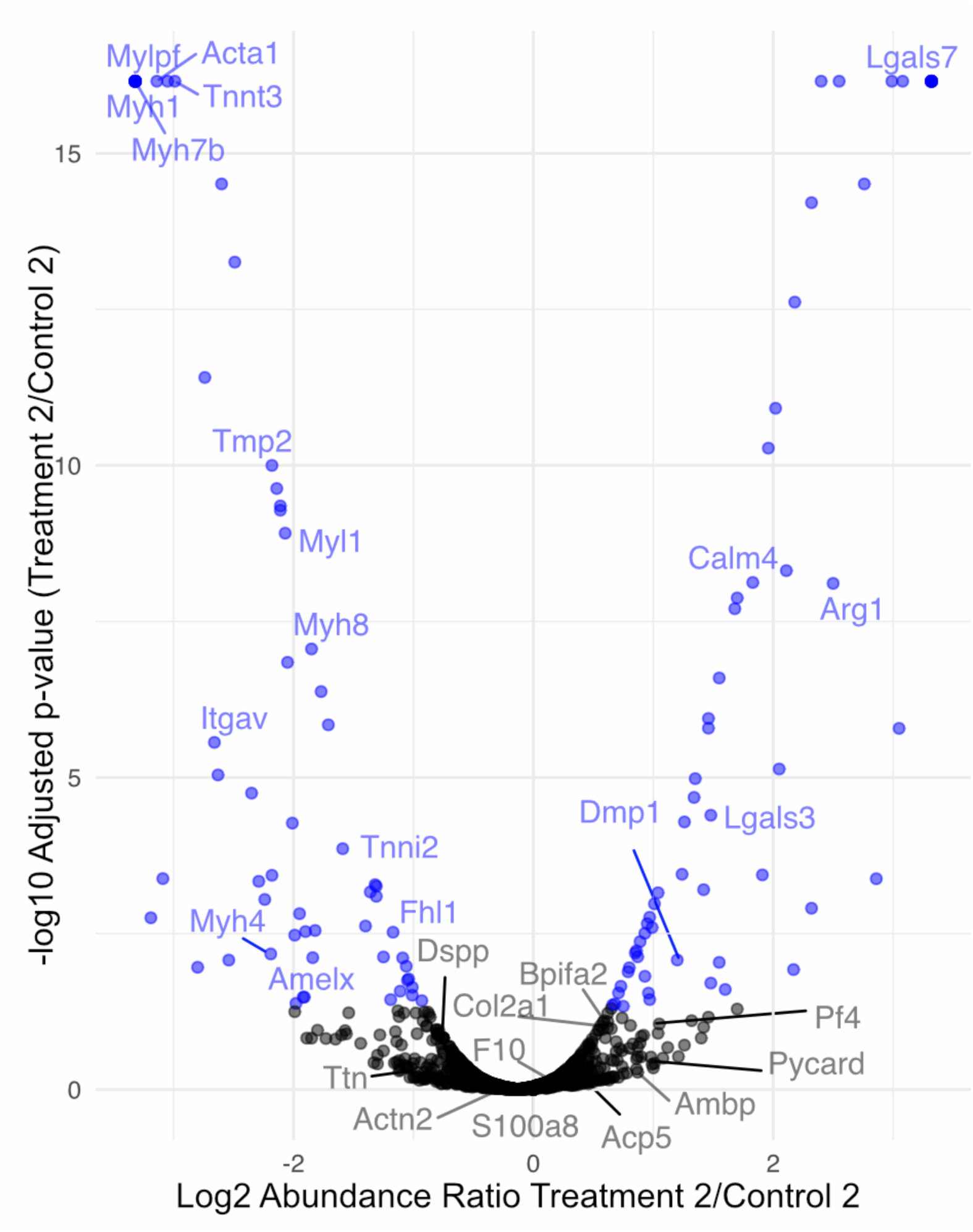
Treatment 2 Over Control 2.

**4:**
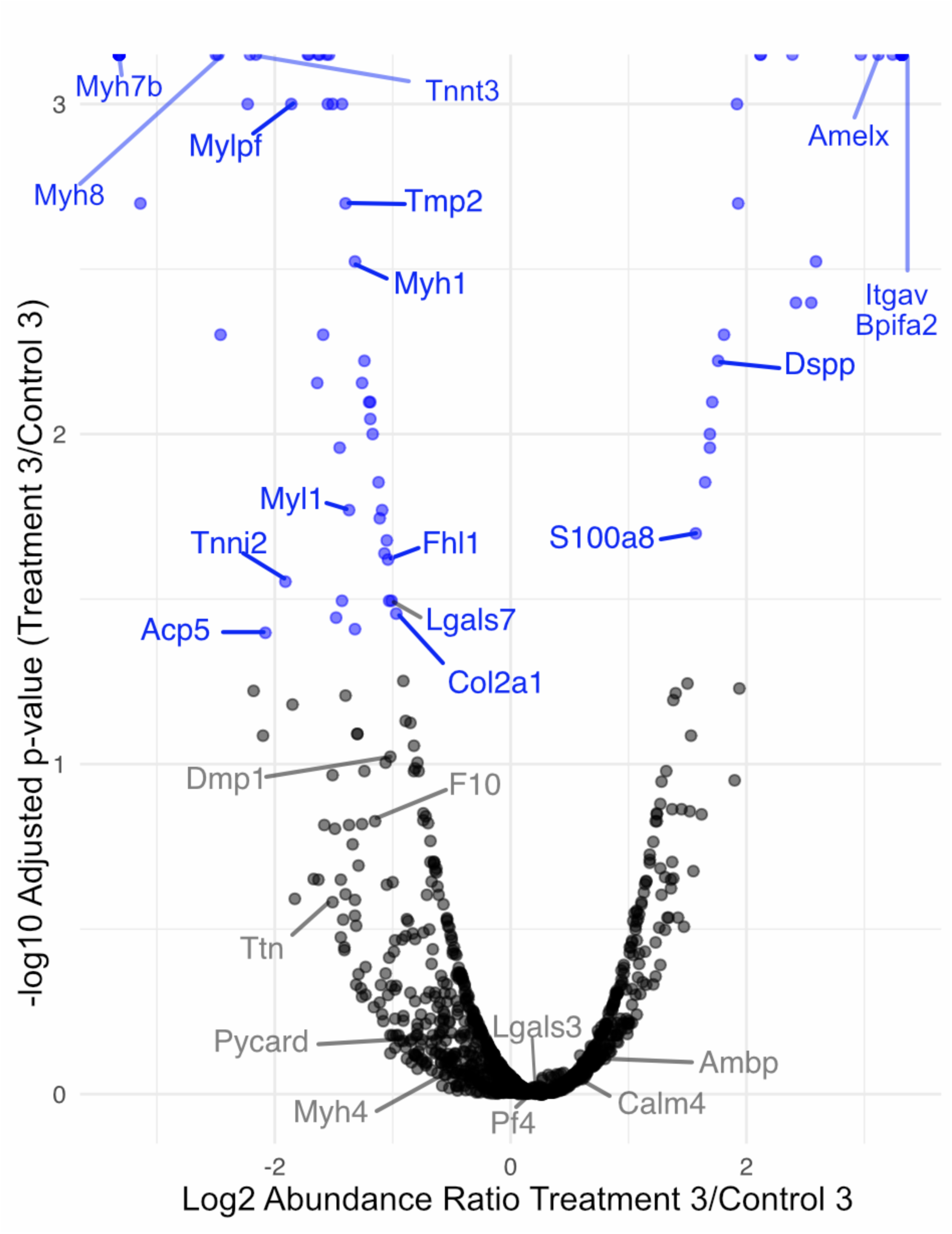
Treatment 3 Over Control 3.

**5:**
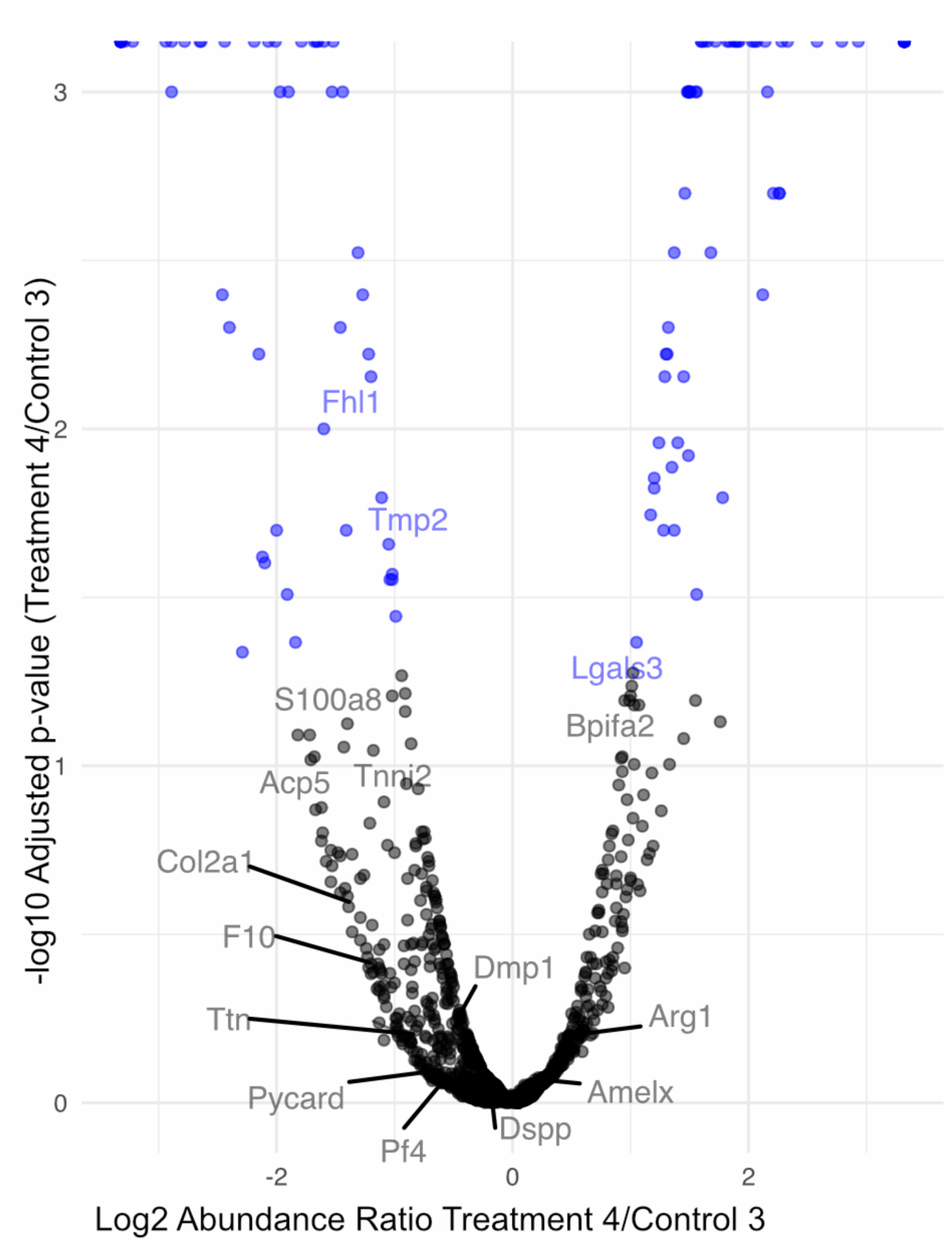
Treatment 4 Over Control 3.

**6:**
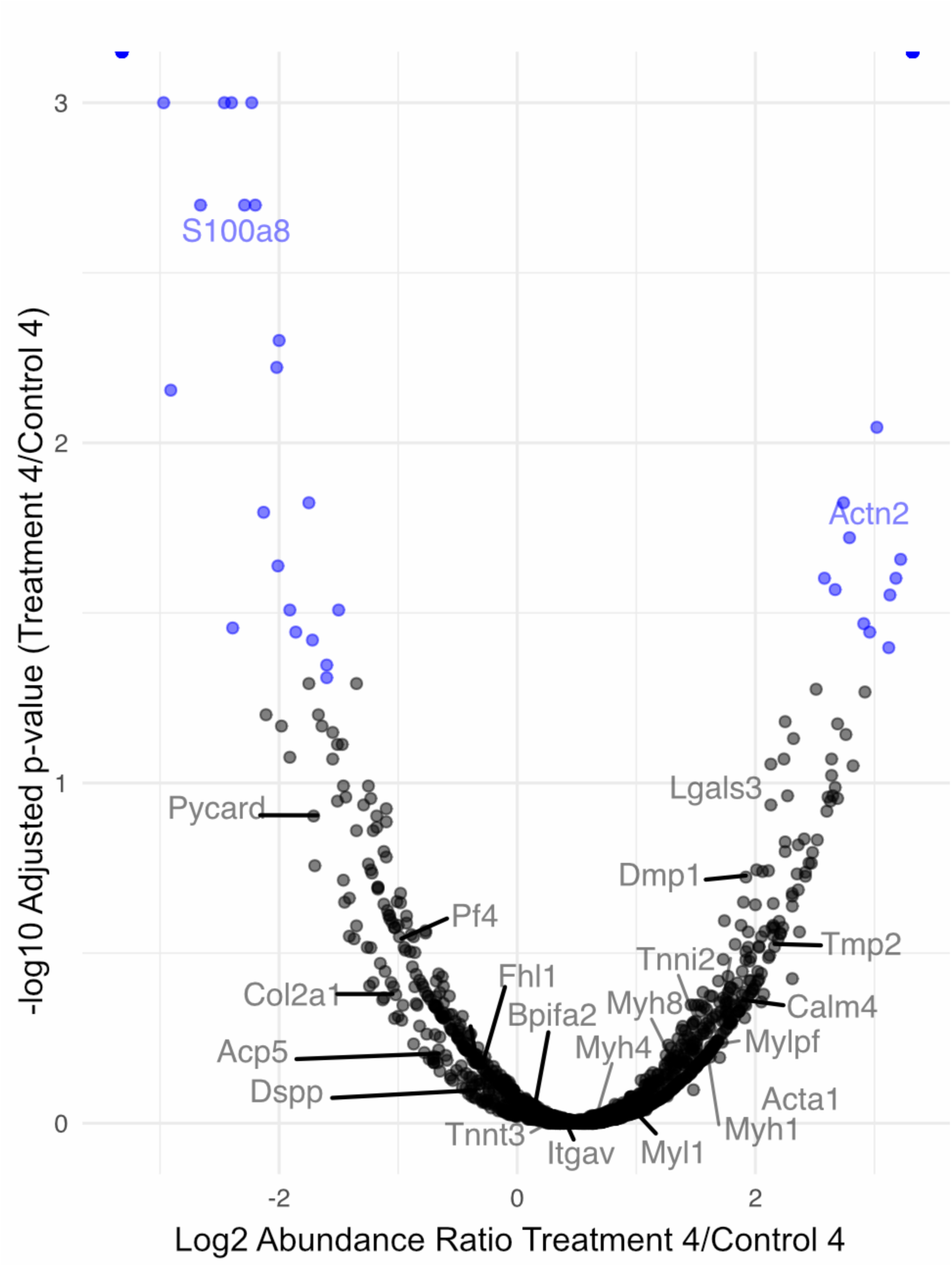
Treatment 4 Over Control 4.

